# Dopaminergic Co-transmission with Sonic Hedgehog Inhibits Abnormal Involuntary Movements

**DOI:** 10.1101/2020.03.09.983759

**Authors:** Lauren Malave, Dustin R. Zuelke, Santiago Uribe-Cano, Lev Starikov, Heike Rebholz, Eitan Friedman, Chuan Qin, Qin Li, Erwan Bezard, Andreas H. Kottmann

## Abstract

L-Dopa induced dyskinesia (LID) is a debilitating side effect of dopamine replacement therapy for Parkinson’s Disease. The mechanistic underpinnings of LID remain obscure. Here we report that diminshed sonic hedgehog (Shh) signaling in the basal ganglia caused by the degeneration of midbrain dopamine neurons (DANs) facilitates the formation and expression of LID. We demonstrate that augmenting Shh signaling with agonists of the Shh effector Smoothened attenuates LID in mouse and macaque models of PD. Employing conditional genetic loss-of-function approaches, we show that reducing Shh secretion from DANs or Smo activity in cholinergic interneurons (CINs) promotes LID. Conversely, the selective expression of constitutively active Smo (SmoM2) in CINs is sufficient to render the sensitized aphakia model of PD resistant to LID. Furthermore, acute depletion of Shh from DANs through prolonged optogenetic stimulation in otherwise intact mice and in the absence of L-Dopa produces LID-like involuntary movements. These findings indicate that augmenting Shh signaling in the L-Dopa treated brain may be a promising and unexpected novel therapeutic approach for mitigating the dyskinetic side effects of long-term treatment with L-Dopa

---

Mesencephalic dopamine neuron (DAN) degeneration results in the hallmark motor and cognitive deficits observed in Parkinson’s disease (PD) [1, 2]. Dopamine (DA) replacement therapy with the DA metabolic precursor L-3,4-dihydroxyphenylalanine (L-Dopa) is the gold-standard treatment for PD and attenuates bradykinesia and akinesia [3, 4]. Unfortunately, prolonged use of L-Dopa therapy leads to a severely debilitating and uncontrollable side effect called L-Dopa induced dyskinesia (LID) that eventually affects ~90 % of all medicated PD patients [5]. These motor impairments impact mobility, produce physical discomfort, negatively impact emotional well-being, and can carry social stigma [6–8].

LID is a medication induced complication that emerges after repeated L-Dopa treatment of PD patients with significant DAN degeneration but does not present in untreated PD or in treated healthy individuals [2, 7, 8]. These observations suggest that DAN degeneration alters brain circuitry such that subsequent DA substitution therapy with L-Dopa is insufficient to fully normalize the functional pathology caused by DAN degeneration and instead promotes LID [9]. While the mechanisms through which DAN degeneration enables LID remain unclear, pathophysiological changes in cholinergic interneurons (CINs) activity have long been suspected to play a critical role in the formation of LID. For example, it is well established that CIN physiology becomes distorted in complex ways following DAN degeneration through changes in morphology, activity, and the post synaptic expression of acetylcholine receptors [10–18]. Additionally, recent work has demonstrated that treatment with L-Dopa alone is not sufficient to restore the normal properties of CINs, as their disturbed activity and morphology remains or becomes further dysregulated following L-Dopa treatment [19, 20]. The complexity of these pathological alterations is highlighted by the fact that both, suppressing and ablating CINs [13, 17, 21], as well as optogenetic stimulation of CINs [22] and pharmacological augmentation of cholinergic signaling can attenuate LID [23].

CINs are direct projection targets of DANs and DA can influence CIN physiology by binding to both inhibitory D2 receptors and potentially facilitatory D5 receptors [24–34]. Dopaminergic control of CINs is further complicated by the fact that DANs also communicate with their targets via the neurotransmitters glutamate [35, 36], GABA [37], and the axonaly transported and activity-dependent secreted peptide sonic hedgehog (Shh) [38–42]. The consideration that DA independent mechanisms of communication between DANs and CINs are present in the healthy brain but absent in the PD brain due to DAN degeneration suggests that LID might arise in part because L-Dopa treatment alone fails to augment additional modalities of dopaminergic co-transmission. Despite this possibility, whether the loss of dopaminergic co-transmitters following DAN degeneration renders brain circuits susceptible to LID remains to be studied.

Unlike glutamate [43–46] and GABA [37], Shh is expressed by all mesencephalic DANs [38]. Post-synaptically, all CINs express the Shh receptor Patched and downstream effector Smoothened (Smo) [38] (Fig. 1A). DAN degeneration in PD or animal models should therefore cause a diminishment in the activity of both DA and Shh signaling pathways among CINs (Fig. 1A). The subsequent administration of L-Dopa would then expose CINs to increased levels of DA but relatively diminished levels of Shh (Fig. 1A). We therefore tested the possibility that LID emerge in part because of the L-Dopa treatment-caused imbalance of Shh and DA signaling onto CINs in the hypodopaminergic brain. More specifically, we hypothesized that augmenting L-Dopa therapy with agonists of Shh signaling could counteract LID induction and expression. Our results confirm these hypotheses.

**Figure 1:**
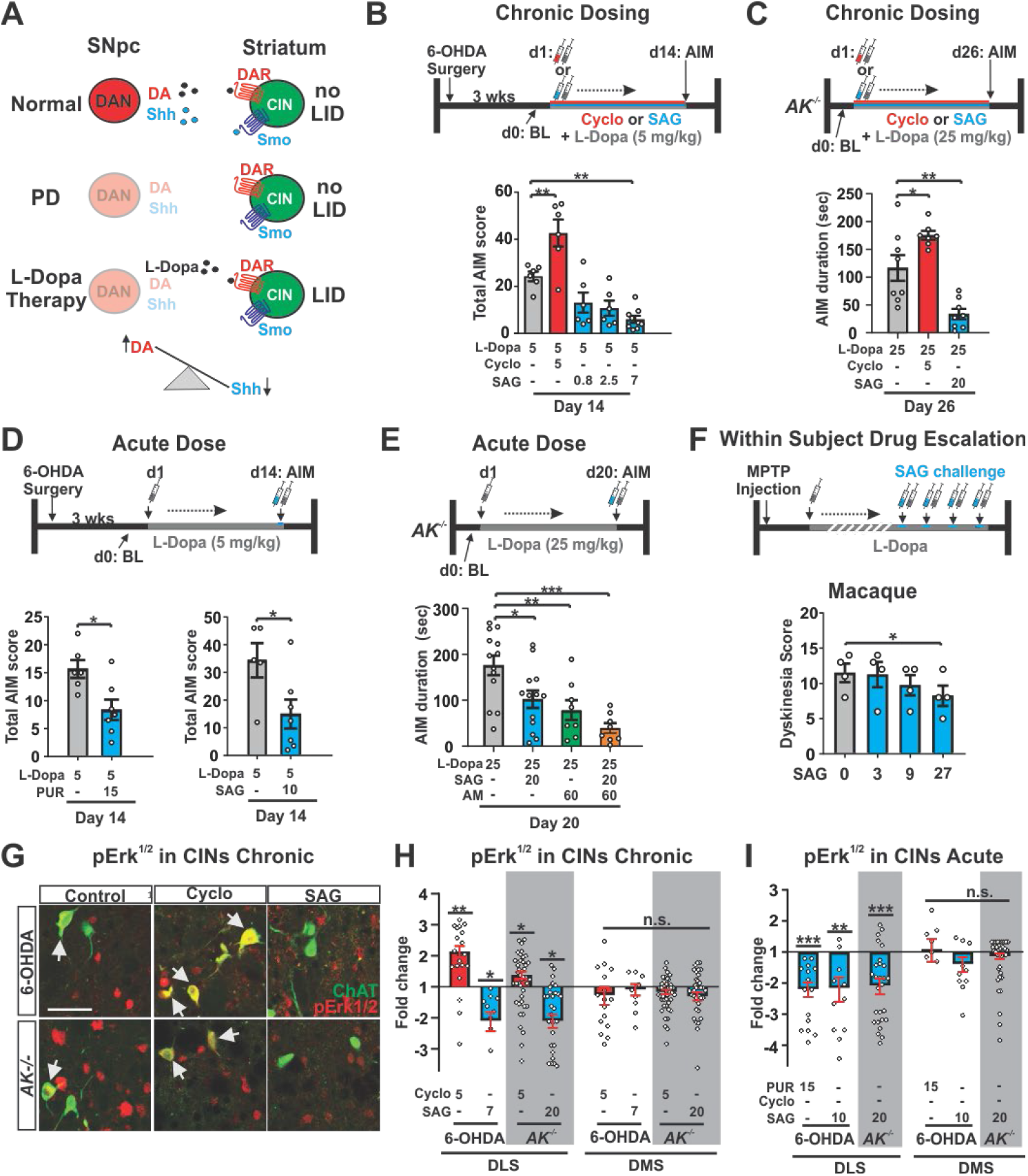
Pharmachological manipulation of Smo bidirectionaly modulates LID in neurotoxic and genetic models of PD. **(A)** DA substitution therapy in the PD brain without augmentation of Shh_DAN_ signaling likely causes an imbalance between DA (high) and Shh (low) signaling onto CINs. **(B)** 6-OHDA mice dosed daily for 14 days with L-Dopa (5 mg/kg) expressed robust LID as shown previously ([48, 75]); white bar). Daily co-injection of L-Dopa with the Smo antagonist Cyclopamine (5 mg/kg, red bar) increased LID while daily co-injection of L-Dopa with the Smo agonist SAG (0.8, 2.5, or 7 mg/kg; blue bars) decreased LID (n= 69; RM two-way ANOVA: F _(4,28)_ = 12.24, p<0.0001; post hoc Bonferroni’s test:* P<0.05, ** P<0.01 treatment vs. control). **(C)** Compared to AIM duration observed in *AK*^-/-^mice dosed daily for 26 days with L-Dopa (25 mg/kg, shown previously to induce robust LID in this paradigm ([53]) and vehicle, daily co-injection of L-Dopa with Cyclo (5 mg/kg; red bar) extended AIMs duration while daily co-injection of L-Dopa with SAG (20 mg/kg; blue bar) decreased AIMs duration (n=7-8; one-way ANOVA: F _(2,20)_ = 20.46, P<0.0001; post hoc Bonferroni’s test: * P<0.05, ** P<0.01 treatment vs. control). **(D)** A single dose of the Smo agonist purmorphamine (PUR; 15 mg/kg; blue bar, left) or SAG (10 mg/kg; blue bar; right) was capable of attenuating LID established across 14 days of daily L-Dopa (5 mg/kg) dosing (n=7 per treatment; unpaired two-tailed student’s t test * P<0.05 treatment vs. control). **(E)** Expression of AIMs established through daily L-Dopa (25 mg/kg) dosing in *AK*^-/-^ mice was reduced after a single-dose treatment of either SAG (20 mg/kg, blue bar), amantadine (AM, 60 mg/kg, grey bar), or SAG combined with AM (black bar) compared to vehicle-treated controls (white bar) (n= 8-13 per treatment; one-way ANOVA, F _(4,50)_ = 9.79, P<0.0001; post hoc Bonferroni’s test:* P<0.05, ** P<0.01, *** P<0.001 treatment vs. control). **(F)** Dyskinesia score in dyskinetic Macaques was reduced in response to a within-subject drug escalation of SAG (0, 3, 9, and 27 mg/kg; n=4; (Friedman’s nonparametric RM one-way ANOVA followed by Dunn’s post hoc test, *P< 0.05). **(G)** Representative images from DLS of 6-OHDA or *AK*^-/-^ mice showing co-localization (white arrows) of the cytohistological LID marker pErk^1/2^ (red) with the CIN marker ChAT (green) after repeated co-administration of L-Dopa (5 mg/kg for 6-OHDA or 25 mg/kg for *AK*^-/-^ animals) with vehicle (control), cyclo (5 mg/kg), or SAG (7 mg/kg for 6-OHDA mice or 20 mg/kg for *AK*^-/-^ mice). (Post mortem analysis of animals whose AIMs were quantified in panels (B) and (C); Scale bar = 50 μm). **(H)** Quantification of the prevalence of pErk^1/2^-positive CINs shown in (G) expressed as fold change over vehicle in the DLS and DMS. Cyclo treatment increased pErk^1/2^ prevalence and SAG treatment decreased pErk^1/2^ prevalence in the DSLwith no changes in the DMS (n= 3-6 per condition; 3 sections each; ~36 CINs per section, post-mortem analysis of animals quantified in panels (B) and (C)); unpaired two-tailed student’s t test: * P<0.05, ** P<0.01 treatment vs. vehicle). **(I)** Quantification of the prevalence of pErk^1/2^-positive CINs in L-Dopa treated 6-OHDA (5 mg/kg L-Dopa for 14 days) or *AK*^-/-^ (25 mg/kg L-Dopa for 20 days) animals given a single dose of PUR (15 mg/kg) or SAG (10 mg/kg for 6-OHDA animals, 20 mg/kg for *AK*^-/-^ animals). Results are expressed as fold change in pErk^1/2^ exspression over vehicle controls in the DLS and DMS. Smo agonist treatment significantly reduced the prevalence of pErk^1/2^-positive CINs in the DLS but DMS (n= 3-6 per condition; 3 sections each; ~36 CINs per section, post-mortem analysis of animals whose AIMs were quantified in panels (D) and (E); unpaired two-tailed student’s t test: ** P<0.01, *** P<0.001, for treatment vs. vehicle. n.s. indicates P>0.05). All bar graphs are plotted as mean +/- SEM.

## Results

### LID is attenuated by Increasing Smoothened Signaling in Three Models of Parkinson’s Disease

We first tested whether augmentation of Shh signaling throughout L-Dopa therapy could attenuate the development of dyskinesia in preclinical models of LID. Behavioral tests with predictive validity for examining LID have been developed in rodent and non-human primate models which rely on genetic or neurotoxin-induced degeneration of DANs. We quantified LID by a subjective measure called the abnormal involuntary movement (AIM) scale which is sensitive to both LID duration and topographic LID subtypes [7, 47, 48]. In the 6-OHDA rodent model, unilateral striatal injections of the dopaminergic toxin 6-OHDA results in dopaminergic denervation of the ipsilateral striatum and a rotational bias of locomotion ([48] and extended data Fig. 1). As previously established [47], daily injections of L-Dopa (5 mg/kg) in these animals led to the appearance of unilateral AIMs (Fig. 1 B, white bar). We observed that repeated co-injection of L-Dopa with Smo antagonist Cyclopamine [49] increased AIM scores (Fig 1 B, red bar). Conversely, repeated co-injection of L-Dopa with three different doses of Smo agonist SAG [50] resulted in a dose-dependent attenuation of AIMs (Fig. 1 B, blue bars).

We next tested this effect in the aphakia (*AK*^-/-^) mouse line where absence of the transcription factor Pitx3 during development results in a bilateral loss of dopaminergic innervation to the dorsal striatum and hypodopaminergia in the adult brain [51, 52]. As previously shown, daily injection of *AK*^-/-^ mice with L-Dopa at 25 mg/kg resulted in LID [53] (Fig. 1 C, white bar). In agreement with our 6-OHDA lesioned mouse results, repeated co-injections of L-Dopa with Cyclopamine (5 mg/kg) increased the duration of AIMs in *AK*^-/-^ mice (Fig. 1 C, red bar), while daily co-injections of L-Dopa with SAG (20 mg/kg) resulted in a threefold attenuation of AIMs (Fig. 1 C, blue bar) compared to littermate controls. A dose response study revealed that the dose of SAG needed to attenuate LID increased linearly with the dose of L-Dopa used to induce dyskinesia (extended data Fig. 2). These findings indicate that the relative imbalance between Shh and DA signaling determines the severity of LID.

Interestingly, we observed that AIMs could be repeatedly attenuated or reinstated, respectively, by dosing the same animals sequentially with or without SAG in addition to L-Dopa (extended data Fig. 3). We therefore tested whether acute Shh pathway stimulation was sufficient to attenuate the expression of established LID. A single dose of Smo agonist administered with the final dose of L-Dopa after repeated daily dosing with L-Dopa alone revealed that acute exposure to Smo agonists Purmorphamine [54] (PUR, 15 mg/kg) or SAG (10 mg/kg) significantly reduced AIMs in the unilateral 6-OHDA model of LID (Fig. 1 D, blue bars). A single dose of SAG (20 mg/kg) in the *AK*^-/-^ mice following repeated L-Dopa administration alone also reduced AIMs to the same degree that was achieved with the anti-dyskinetic bench mark drug amantadine (AM, 60 mg/kg) [55] (Fig. 1 E, blue and gray bars). Unlike AM, however, neither acute nor chronic dosing with SAG reduced the anti-akinetic benefit of L-Dopa in either of the murine models tested (extended data Fig. 4 B, C). Further, combining SAG (20 mg/kg) and AM (60 mg/kg) produced an additive effect on AIM attenuation suggesting distinct mechanisms of LID attenuation (Fig. 1 E, black bar).

In macaques, treatment with the toxin 1-methyl-4-phenyl-1,2,3,6-tetrahydropyridine (MPTP) produces PD-like DAN degeneration and allows the establishment of LID following chronic L-Dopa that can be attenuated by AM [56]. We utilized individually-tailored L-Dopa doses (see Methods) in this model of LID to further validate the efficacy of acute Smo activation on LID attenuation. Parkinsonian macaques treated with a single co-injection of SAG at 27 mg/kg (the highest dose used in a within subject dose escalation experiment) alongside L-Dopa showed reduced dyskinesia during a two hour observation period (Fig. 1 F). These results suggest that acute activation of the Shh/Smo pathway in non-human primate models of LID is capable of attenuating AIMs to the same degree as is achieved by AM and, as observed in the murine models, is not detrimental to the anti-akinetic effect of L-Dopa (extended data Fig. 4).

Phosphorylation of extracellular signal-regulated protein kinase (Erk) in the striatum is a cell physiological marker for LID [57]. During early LID, pErk^1/2^ (phospho-Thr202/Tyr204– ERK^1/2^) is observed in a dispersed pattern within the striatum among primarily medium spiny projection neurons (MSN) [57]. However, dosing with L-Dopa progressively increases pErk^1/2^ prevalence among CINs in the dorsolateral striatum (DLS) in correlation with increased LID severity [13]. Conversely, pharmacological inhibition of Erk phosphorylation attenuates LID [13]. We found that in 6-OHDA and *AK*^-/-^ mice, daily co-injections of L-Dopa and inhibitory (Cyclopamine; Fig. 1 H, red bars) or stimulatory (SAG; Fig. 1 H, blue bars) Smo pharmacology resulted in a respective increase or decrease in the fraction of pErk^1/2^-positive CINs of the DLS 30 min after the final co-administration (Fig. 1 G, H). The fraction of pErk^1/2^-positive CINs in the dorsomedial striatum (DMS) of the same animals was not affected. Similarly, and concordant with the respective behavioral results, a single dose of the Smo agonists SAG or PUR co-injected with the final daily dose of L-Dopa reduced the prevalence of pErk^1/2^ positive CINs in the DLS but not DMS of both 6-OHDA and *AK*^-/-^ mice (Fig. 1 I).

Together, these results from mouse and macaque models of LID indicate that Smo activity modulates the formation and expression of AIMs as well as the activation of LID marker pErk in CINs of the DLS through a reversible, bi-directional, immediate, and dose-dependent mechanism.

### Conditional Ablation of Smo from CINs facilitates LID while expression of constitutively active SmoM2 in CINs of *AK*^-/-^ mice blocks LID

Given the bidirectional changes in Erk phosphorylation we observed among CINs in response to Smo pharmacology and previous findings that CINs express the Shh receptor Patched 1 [38], we were led to the possibility that Shh signaling impinges on LID through CINs. If Smo expressed by CINs is in fact the relevant target of Smo antagonists and agonists that respectively promote or attenuate LID and Erk activation, then the conditional, genetic loss or gain of function of Smo in CINs should phenocopy the pharmacological outcomes. Confirming this expectation, we found that selective ablation of Smo from CINs (*Smo_ChAT-Cre^-/-^_*, Fig. 2 A) in otherwise intact mice receiving daily L-Dopa dosing (25 mg/kg, Fig. 2 B) facilitated LID compared to heterozygous littermate controls (Fig. 2 C). Concordant with the behavior, these mice also exhibited a corresponding increase in the fraction of pErk^1/2^-positive CINs of the DLS (extended data Fig. 5 A). Conversely, expression of constitutively active Smo (the oncogene SmoM2 [58]) selectively in CINs of *AK*^-/-^ mice (*SmoM2_ChAT-Cre_^+/-^; AK^-/-^*, Fig. 2 D, extended data Fig. 6) that received daily L-Dopa dosing (25 mg/kg, Fig. 2 E) blocked the formation and progressive intensification of AIMs observed readily in *AK*^-/-^ littermate controls (Fig. 2 F). Concordant with this resistance to LID formation, *SmoM2_ChAT-Cre_*^+/-^; AK^-/-^ mice displayed a corresponding decrease in the prevalence of pErk^1/2^-positive CINs of the DLS compared to *AK*^-/-^ control mice (extended data Fig. 5 B).

**Figure 2:**
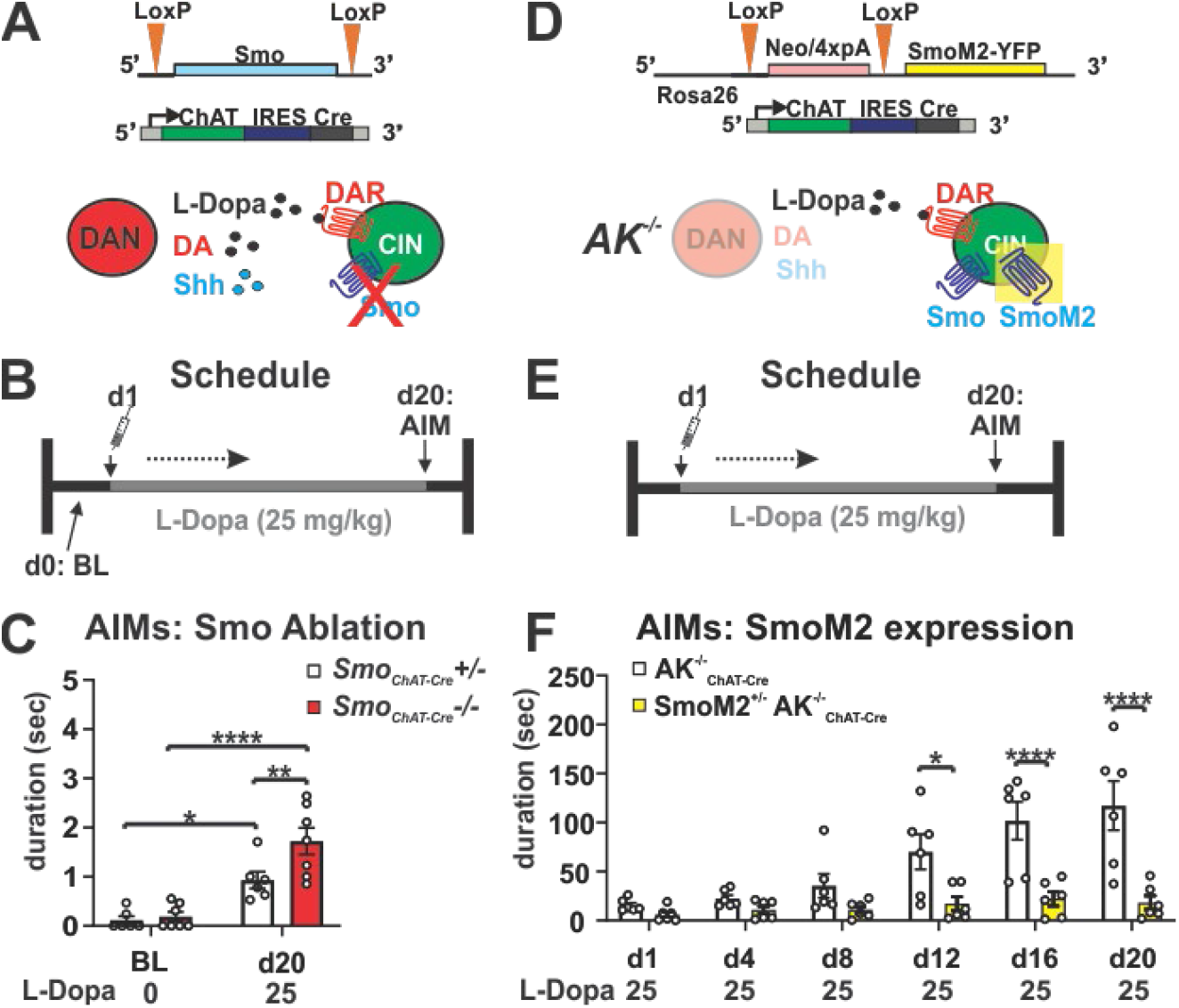
Conditional genetic ablation of Smo from CINfacilitates LID while conditional expression of constitutive active SmoM2 in CIN inhibits LID. **(A)** Smo was selectively ablated from cholinergic neurons by ChAT-IRESCre, which renders CINs insensitive to endogenous Shh signaling. **(B)** L-Dopa dosing schedule for the Smo loss-of-function study. **(C)** AIMs are increased in *Smo_ChAT-Cre_*^-/-^ mice compared to *Smo_ChAT-Cre_*^-/+^ heterozygous control littermates in response to daily injection of 25 mg/kg L-Dopa for 20 days (n = 6-7 per genotype; RM two-way ANOVA day x genotype effect: F _(1,11)_ 4.95, p=0.048. Post hoc Bonferroni’s test: ** P<0.01 control vs. mutant on d20; day effect: F _(1,11)_ = 52.61, p<0.0001; * P<0.05 control BL vs. d20 and **** P<0.0001 mutant BL vs. d20). **(D)** Constitutively active SmoM2 was selectively expressed in cholinergic neurons of *AK*^-/-^ mice by ChAT-IRESCre, thus activating Smo signaling in CIN independent of endogenous Shh availibility. **(E)** L-Dopa dosing schedule for the SmoM2 gain-of-function study. **(F)** AIMs formation in response to daily injection of 25 mg/kg L-Dopa is suppressed in *SmoM2*^+/-^ *AK^-/-^_ChAT-Cre_* mice compared to *AK*^-/-^_*Chat-Cre*_ littermate controls (n = 6 per genotype; RM two-way ANOVA time x genotype effect: F _(5,50)_ =8.16, p=<0.0001. Post hoc Bonferroni’s test: * P<0.05, **** P<0.0001 control vs. mutant).

### Shh Originating from DANs Suppresses Abnormal Involuntary Movements

Given that DANs of the Substantia Nigra pars compacta (SNpc) express Shh (Shh_DAN_) throughout life, target CINs of the DLS, and degenerate in PD, these cells are a likely physiological source of Shh that could impinge on Smo activity in CINs [38, 59, 60]. In accordance with the idea that Shh_DAN_ expression impacts LID, we found that mice with conditional ablation of Shh from DANs (*Shh_DAN_^-/-^* [38] Fig. 3 A) displayed AIMs in response to daily L-Dopa dosing (Fig. 3 B, C) in a dose-dependent manner (d11 vs d12) while heterozygous littermate controls did not. To determine if post synaptic activation of Smo by SAG could rescue the LID phenotype of *Shh_DAN_*^-/-^ mice, we injected *Shh_DAN_*^-/-^ and heterozygous control mice with 25 mg/kg L-Dopa daily for 20 days followed by either 10 or 20 mg/kg SAG alongside the final L-Dopa dose (Fig. 3 D). As before (Fig. 3 C), we found that L-Dopa dosing resulted in AIMs among *Shh_DAN_*^-/-^ mice but not heterozygous littermate controls (Fig. 3 E). A single dose of SAG was able to significantly attenuate LID at both concentrations in *Shh_DAN_^-/-^* mice, with the higher concentration having a more profound effect (Fig. 3 E). Concordant with the behavioral phenotype, L-Dopa dosing in *Shh_DAN_*^-/-^ mice resulted in an increase in the fraction of pErk^1/2^-positive CINs of the DLS compared to controls, which acute SAG co-injection with L-Dopa normalized to control levels (Fig. 3 F, G). As observed in the 6-OHDA and *AK*^-/-^ models, L-Dopa dosing in *Shh_DAN_*^-/-^ mice did not affect the prevalence of pErk^1,2^-positive CINs in the DMS (Fig. 3 G).

**Figure 3:**
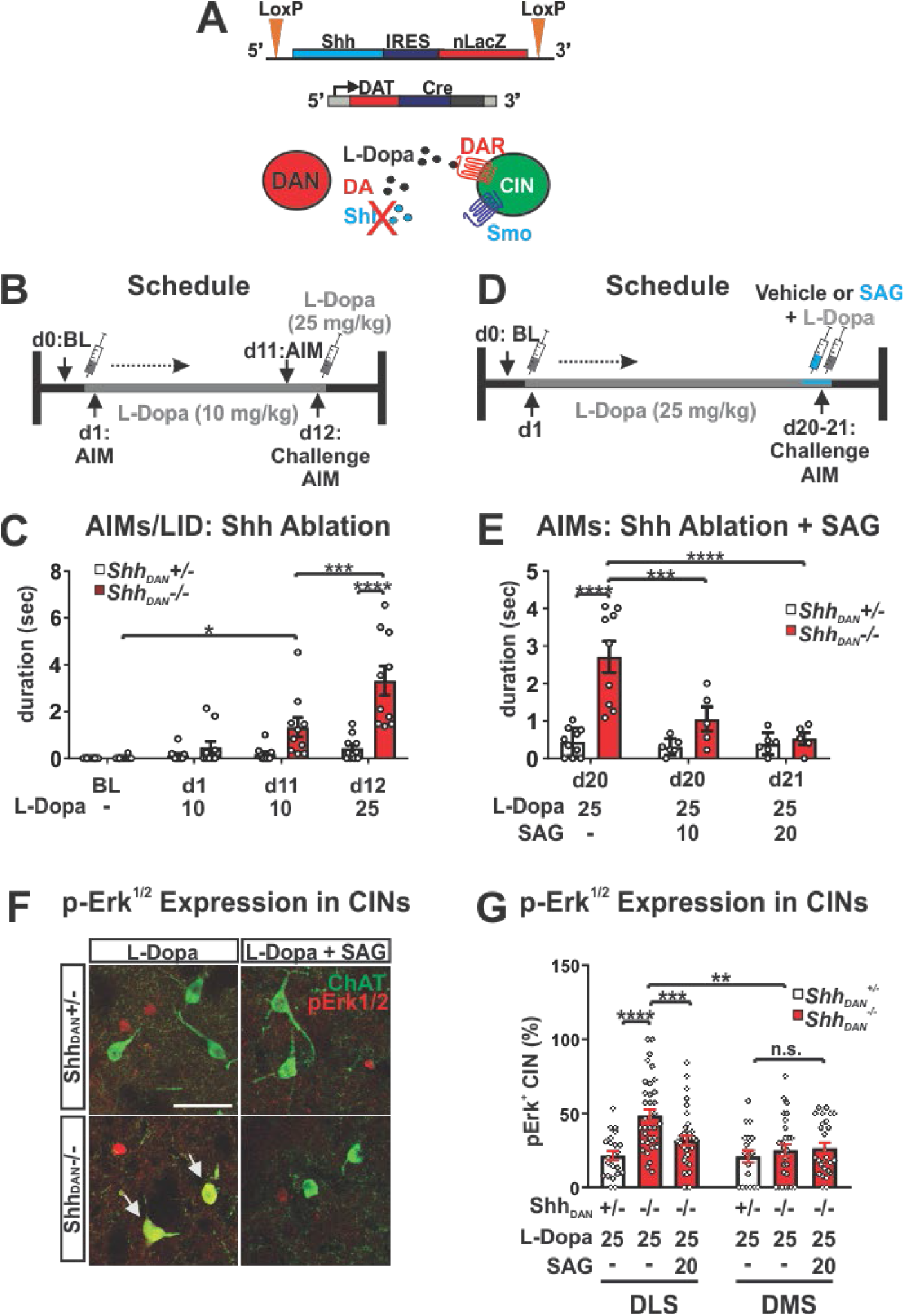
Conditional genetic ablation of Shh from DAN facilitates AIMs which can be attenuated by agonists of Smo. **(A)** Shh was selectively ablated from DANs by Dat-Cre which reduces Shh/Smo signaling in the striatum [38]. **(B)** L-Dopa dosing schedule for the Shh_DAN_ loss-of-function study. **(C)** Daily injection of 10 mg/kg L-Dopa for 11 days in *Shh_DAN_*^-/-^ mice elicited AIMs which were further increased in response to a 25 mg/kg L-Dopa challenge dose. This effect was absent in *Shh_DAN_*^-/+^ heterozygous littermate control mice undergoing the same treatment (n = 10 per genotype; RM two-way ANOVA day x genotype: F _(2,36)_ = 4.53, p=0.018. Post hoc Bonferroni’s test: * P<0.05 for BL vs. d11); treatment effect: F _(1,18)_ = 16.23, p=0.001; post hoc Bonferroni’s test: *** P<0.001 d11 vs. d12; genotype effect: F _(1,18)_ = 18.36, p=0.0004; post hoc Bonferroni’s test: **** P<0.0001 control vs. mutant on d12). **(D)** L-Dopa and SAG dosing schedule for the pharmacological complementation of the LID phenotype seen in *Shh_DAN_*^-/-^ mutants. **(E)** A single injection of SAG (10 mg/kg or 20 mg/kg) attenuated AIMs that were established by daily injections of L-Dopa (25 mg/kg) for 20 days in *Shh_DAN_*^-/-^ mice. No effect was observed in *Shh_DAN_*^+/-^ heterozygous littermate control mice (n = 5–10; two-way ANOVA genotype effect: F _(1,35)_ = 21.36, p=0.0002. Post hoc Bonferroni’s test: **** P<0.0001 control vs. mutant on d 20; genotype x treatment effect: F_(2,35)_=9.15, =0.001. Post hoc Bonferroni’s test:*** P<0.001, **** P<0.0001 mutant with and without SAG on d 20 and d 21). **(F)** Images of pErk^1/2^ (red) co-localization (arrow) with ChAT (green) in the DLS of *Shh_DAN_*^-/-^ mutants and *Shh_DAN_*^-/+^ heterozygous littermate controlsupon chronic daily dosing with L-Dopa alone or L-Dopa with SAG (post mortem analysis of animals shown in (L); scale bar = 50 μm). **(G)** L-Dopa increased the prevalence of pErk^1/2^-positive CINs in the DLS but not DMS of *Shh_DAN_*^-/-^ mice. This increase could be attenuated through co-injection of SAG with L-Dopa (n = 8–12, 35–38 CINs per genotype and drug condition; three-way ANOVA genotype effect: F _(1,105)_ = 7.093, p=0.01; post hoc Bonferroni’s test: **** P<0.0001 control vs. mutant; treatment effect: F _(1,105)_ = 9.79, p=0.002; post hoc Bonferroni’s test: **** P<0.0001 mutant with and without SAG; region x genotype: F _(1,98)_ = 11.21, p=0.001; post hoc Bonferroni’s test: **P<0.01 mutant DLS vs. DMS). All bar graphs are plotted as mean +/- SEM.

These conditional genetic manipulations indicate that Shh_DAN_ to Smo_CIN_ signaling interferes with LID through a mechanism that resides at the level of regulating CIN physiology, and therefore upstream of cholinergic signaling in the striatum.The action of Shh_DAN_ on LID expression via Smo_CIN_ is thus likely antecedent to LID inhibition by muscarinic signaling on D1 MSN [23], or nicotinic receptor desensitization proposed to occur upon optogenetic CIN activation [22] (extended data Fig. 7 A). Concordant, we found that cyclopamine induced intensification of AIMs in *AK*^-/-^ mice (Fig. 1 C, extended data Fig. 7 B) could be attenuated by an acute dose of M4PAM (extended data Fig. 7 C).

Taken together, the results from our conditional pre- and post-synaptic loss of function mouse lines, in combination with our post synaptic gain of function paradigm, indicate that interruption of Shh signaling from DANs to CINs of the DLS promotes LID formation and Erk activation whereas constitutively active Smo in CINs blocks LID formation and Erk activation in the otherwise LID sensitive *AK*^-/-^ model. The Shh_DAN_/Smo_CIN_ dependent mechanisms highlighted by these paradigms appears to function upstream of changes in striatal cholinergic signaling and independent of DAN projection degeneration or maintenance in the striatum (extended data Fig. 8).

### Depletion of Shh Signaling from DANs via Prolonged Stimulation Facilitates Abnormal Involuntary Movements

The changes in DAN co-transmission probed by our pharmacological and genetic experiments relied on neurotoxic and genetic LID models with long term reductions in Shh_DAN_ and therefore likely functional and structural compensatory changes to basal ganglia physiology. However, the observation that LID could be attenuated through acute Smo pharmacology raised into question the assumption that long-term reduction of Shh_DAN_ and possible compensatory adaptation is required for LID-like pathology. In order to resolve whether long-term adaptations following a reduction in Shh_DAN_ signaling are a prerequisite for LID, we sought to develop a paradigm in which Shh_DAN_ depletion could be achieved within minutes in an otherwise intact brain.

High-frequency stimulation is capable of triggering Shh release from neurons [42]. However, prolonged stimulation of neurons can produce an acute exhaustion of peptide stores and deplete peptide signaling over time in a manner not observed with small neurotransmitters [61–63]. We therefore explored whether imposing prolonged dopaminergic burst activity through optogenetic stimulation would result in a progressive reduction of Shh_DAN_ signaling. Forced, unilateral pulsatile burst firing of DANs (Fig. 4 A, [42]) resulted in optical stimulation-dependent rotational activity that intensified over the course of an hour of stimulation, and thus indicated a linear change in DAN signaling across the session (Fig. 4 B). These effects were observed without detectable change in post-mortem dopaminergic fiber density within the striatum (extended data Fig. 8 and 9). To determine whether the intensification of rotational activity was in part elicited by the exhaustion of Shh signaling, we examined the effects of Smo pharmacology on rotational behavior. Smo Inhibition via Cyclopamine injection (5mg/kg) 30 min prior to onset of optogenetic stimulation elicited a maximal rotational response immediately at the beginning of stimulation, which otherwise was only observed after one hour of prolonged stimulation in carrier injected control mice (Fig. 4 C). Conversely, injection of the Smo agonist SAG (20 mg/kg) 30 min prior to onset of optogenetic stimulation blocked the gradual increase in circling behavior seen in control mice (Fig. 4 D). Thus, the bidirectional effects of Smo agonist and antagonist treatment on the progressive intensification of circling behavior indicated that optogenetic stimulation of DANs produced gradual diminshment of Shh_DAN_ signaling. Given the differences in susceptibility to transmitter exhaustion between DA and Shh, these results suggest that prolonged forced stimulation of DANs creates an imbalance of relatively high DA and diminished Shh_DAN_ signaling onto CINs, akin to that which facilitates the expression of AIMs (Fig. 1 – 3).

**Figure 4:**
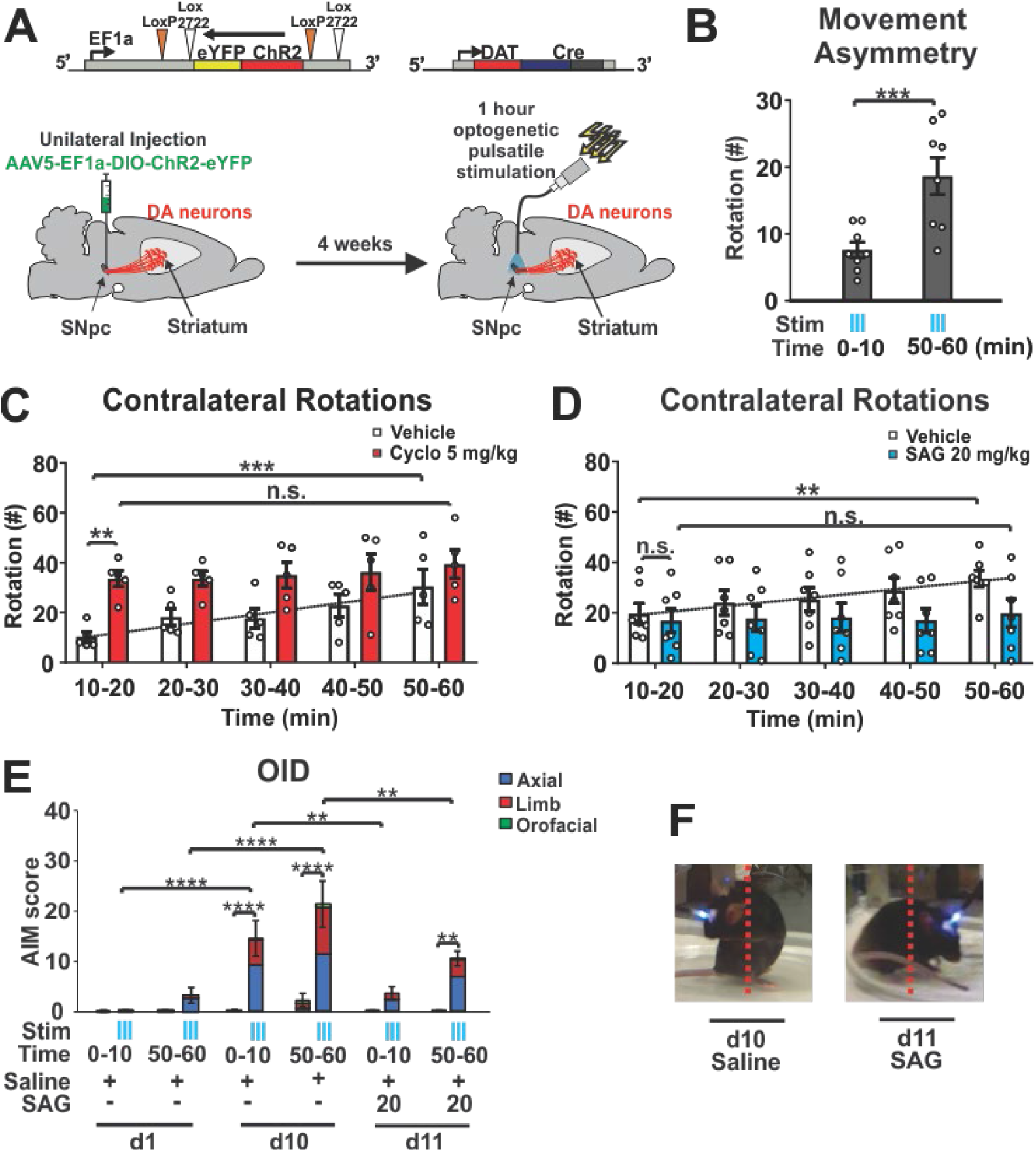
Prolonged optogenetic stimulation of DANs depletes Shh_DAN_ signaling and facilitates AIM-like behavior. **(A)** Channelrhodopsin (ChR2) was expressed unilaterally in DANs. Following 4 weeks recovery period, DANs were stimulated daily across an hour session in 5 second-long bouts separated by 30 second pauses. Bouts of optogenetic stimulation consisted of 10-msec light pulses at 60 Hz. AIMs were scored throughout the hour-long session. **(B)** Stimulation resulted in a contralateral rotational bias which increased across each hour long session (n = 8; paired two-tailed Student’s t test: ***P<0.001). **(C)** Number of contralateral rotations in 10 min bins over the course of an hour-long stimulation session in animals injected with vehicle (white bars) or Cyclopamine (5 mg/kg; red bars; n = 5 per condition). RM two-way ANOVA treatment effect: F_(1,8)_= 7.40 p=0.026. Post hoc Bonferroni’s test: **P<0.01 vehicle vs. cyclopamine (Cyclo); time effect F_(4,32)_= 5.69 p=0.001. Post hoc Bonferroni’s test: ***P<0.001 veh 10-20 min vs. veh 50-60 min. **(D)** Contralateral rotations as in (C) following vehicle or SAG (20 mg/kg; n = 7 per condition; RM two-way ANOVA, time effect: F_(4,44)_= 6.96 p=0.0002; post hoc Bonferroni’s test: **P<0.01 veh 10–20 min vs. veh 50–60 min). **(E)** Optogeneteically induced dyskinesia comprised of axial (red), limb (green), and orofacial (yellow) AIMs were scored in the absence and presence of stimulation (blue ticks) during the first and last 10 mins of each stimulation session. Results are reported for days 1, 10, and 11, the last of which, SAG (20mg/kg) was administered prior to the start of stimulation (n = 8; two-way ANOVA, stimulation effect: F _(1,38)_ = 61.1, p<0.0001; post hoc Bonferroni’s test: **** P<0.0001 stimulation vs. non-stimulation d10 and **P<0.01 stimulation vs. non-stimulation d11; time x stimulation effect: F _(15,38)_ = 8.59, p<0.0001; post hoc Bonferroni’s test: **** P<0.0001 stimulation during d1 vs. d10 and **P<0.01 stimulation during day10 vs. d11). **(F)** Still video images of mouse posture during stimulation with and without SAG (20 mg/kg) injection (representative videos in supplemental files).

Concordant, we found that LID-like axial and limb AIMs, which we identified as optical induced dyskinesias (OID), emerged towards the end of the stimulation session on day 1 and were time-locked with laser onset (Fig. 4 E AIMs summed in 10 minute bins, extended data video OID day 1 control). These dyskinetic movements intensified across daily one-hour stimulation sessions. On day 10, we observed orofacial movements (tongue protrusions) in addition to abnormal axial and limb movements which emerged during the first 10 minutes of stimulation and intensified throughout the session (Fig. 4 E, extended data video OID day 10). A single dose of SAG administered prior to optogenetic stimulation on day 11 was sufficient to significantly attenuate axial, limb, and orofacial OID (Fig. 4 E and F; extended data video OID day 11). These results therefore indicate that LID-like AIMs do not require long-term adaptations following the loss of Shh_DAN_ but can in fact emerge within minutes of pulsatile DAN burst firing and concomitant exhaustion of Shh_DAN_ release.

### Shh Signaling from DANs Elevates Phosphorylation of the Neuronal Activity Marker p-rpS6 in CINs via Smo Activation

The rapid modulation of LID-like behavior following changes in Shh signaling suggested that Shh_DAN_ might impinge on CIN electrical activity via Smo. In CINs, levels of the ribosomal activity marker p-rpS6^240/244^ have been shown to correlate positively with CIN activity under basal and stimulated conditions [64–66]. Therefore, we next examined whether Shh_DAN_/Smo_CIN_ signaling can affect p-rpS6^240/244^ levels averaged across CINs of the DSL.

In agreement with the previous finding that muscarinic autoreceptor M2 expression was increased and, conversely, cholinergic tone reduced in the striatum of *Shh_DAN_*^-/-^ mice [38], we found reduced p-rpS6^240/244^ levels in CINs of *Shh_DAN_*^-/-^ mice compared to controls (Fig 5 A, B). A single dose of SAG administered 30 min prior to analysis, restored p-rpS6^240/244^ levels in *Shh_DAN_*^-/-^ mice (Fig. 5 A, B). The selective ablation of Smo from CINs in *Smo_ChAT-Cre_*^-/-^ animals phenocopied the reduction in p-rpS6^240/244^ levels in CINs of *Shh_DAN_*^-/-^ mice (Fig. 5 C, D). However, SAG failed to restore p-rpS6^240/244^ levels in CINs of *Smo_ChAT-Cre_*^-/-^ animals, indicating that SAG dependent alterations of CIN physiology are direct and that Shh_DAN_ normalizes p-rpS6^240/244^ levels through Smo activation on CINs (Fig. 5 C, D). Mice expressing constitutively active SmoM2 in CINs (*SmoM2_ChAT-Cre_*^+/-^mice) did not exhibit further increases in average p-rpS6^240/244^ levels among CINs, suggesting that basal levels of Shh signaling produce a maximal effect on p-rpS6^240/244^ levels in CINs (Fig. 5 C, D). Furthermore, one hour of DAN optogenetic stimulation decreased p-rpS6^240/244^ levels in CINs of the stimulated ipsilateral DLS, but not in CINs of the unstimulated contralateral side (Fig. 5 E, F). SAG administration prior to onset of optogenetic stimulation blocked this decrease (Fig. 5 E, F). These observations indicate that Shh_DAN_ to SmoCIN signaling contributes to cholinergic activity in a direct and acute manner as measured by p-rpS6^240/244^.

**Figure 5:**
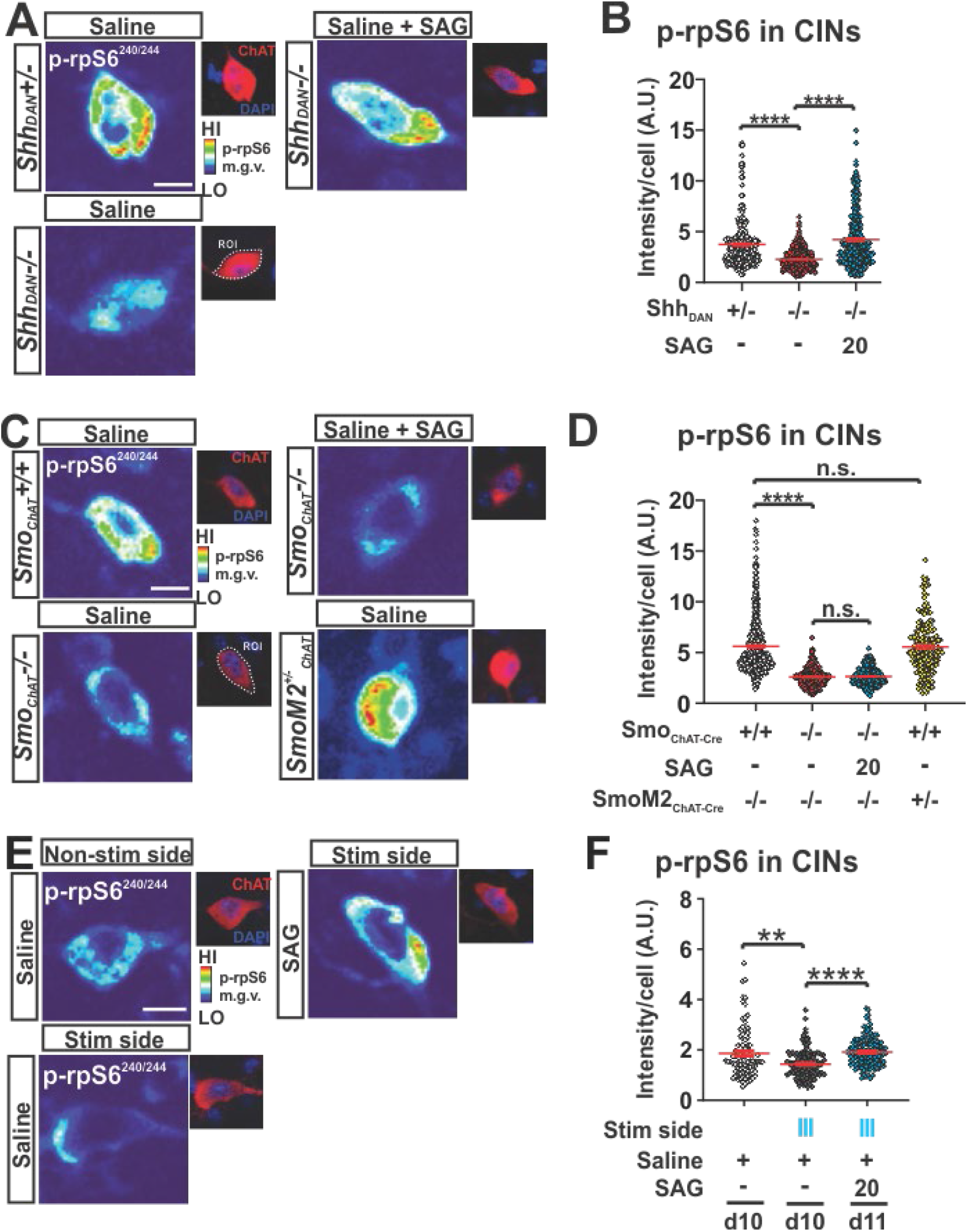
Shh_DAN_ signaling impinges on the physiology of CINs via Smo_CIN_. **(A)** Representative heat maps showing DLS CIN levels of the ribosomal activity marker p-rpS6^240/244^ in *Shh_DAN_*^-/-^ mice, *Shh_DAN_*^-/-^ littermate controls, and in response to a single dose of of SAG (20 mg/kg; p-rpS6^240/244^ levels were normalized to NeuN levels in CINs identified by ChAT staining, see smaller images; Scale bar = 10 μm). **(B)** Quantification of images referenced in (A). Analysis showed reduced average levels of p-rpS6^240/244^ in *Shh_DAN_*^-/-^ mice compared to *SHH_DAN_*^-/+^ littermate controls which were normalized by a single dose of SAG (20 mg/kg) (n = 3–4 per genotype and drug condition, 60–100 cells per n; each dot represents one neuron; red lines indicate the mean ± SEM; Kruskal-Wallis nonparametric one-way ANOVA followed by Dunn’s post-test: **** P<0.0001 controls vs. mutants, or mutants with and without treatment). **(C)** Representative heat maps showing DLS CIN levels of p-rpS6^240/244^ in *Smo^+/+^_ChAT-cre_* mice (loss of function), *SmoM2^+/-^_ChAT-cre_* mice (gain of function), *Smo*^+/+^_*ChAT-cre*_ littermate controls, and in *Smo*^-/-^_*ChAT-cre*_ mice administered SAG (20 mg/kg; Scale bar = 10 μm). **(D)** Quantification of images referenced in (C). Analysis showed reduced average levels of p-rpS6^240/244^ in *Smo*^-/-^_*ChAT-cre*_ mice compared to *Smo*^+/+^_*ChAT-cre*_ littermate controls which could not be normalized by SAG treatment. P-rpS6 levels in *SmoM2*^+/-^_*ChAT-cre*_ trended higher compared to *Smo*^+/+^_*ChAT-cre*_ littermate controls but did not reach significance (n = 3–4 per genotype and treatment group, 60–100 cells per n; Kruskal-Wallis nonparametric one-way ANOVA followed by Dunn’s post-test: **** P<0.0001 controls vs. mutants, or mutants with and without SAG, n.s. indicates P>0.05). (**E)** Representative heat maps showing DLS CIN levels of p-rpS6^240/244^ in response to optogenetic stimulation of DANs (Scale bar = 10 μm). **(F)** Quantification of images referenced in (E). Analysis showed reduced average levels of p-rpS6^240/244^ in CINs of the DLS after one-hour of pulsatile DAN stimulation compared to non stimulated controls also expressing ChR2 in DANs. This reduction was elevated to control levels by a single dose of SAG (20 mg/kg) injected prior to optogenetic stimulation (n = 2 per saline condition and n = 6 for SAG; 40 to 100 cells per n; paired two-tailed Student’s t test: ** P<0.01 stimulated (stim) vs. unstimulated side; unpaired two-tailed Student’s t test: ****P<0.0001 saline vs. SAG).

## Discussion

LID develops in response to pulsatile administration of L-Dopa to the hypodopaminergic brain. Here, we present several lines of evidence showing both that reduced Shh signaling from DANs to CINs facilitates the formation and expression of LID, and that augmenting Shh signaling attenuates LID. We observed that: (1) In the 6-OHDA mouse, MPTP macaque, and *AK*^-/-^ genetic mouse models of DAN loss, agonists of the Shh signaling effector Smo attenuate LID formation and the expression of established LID (Fig. 6A). (2) Pre- and post-synaptic conditional ablation of Shh from DANs or Smo from CINs, respectively, facilitates LID (Fig. 6B). (3) Cholinergic neuron-specific expression of constitutively active Smo, SmoM2, blocks LID formation in the *AK*^-/-^ mouse model of LID (Fig. 6C). (4) Acute depletion of Shh_DAN_ by prolonged optogenetic stimulation of DANs in otherwise intact mice results in LID-like AIMs (Fig. 6D).

**Figure 6:**
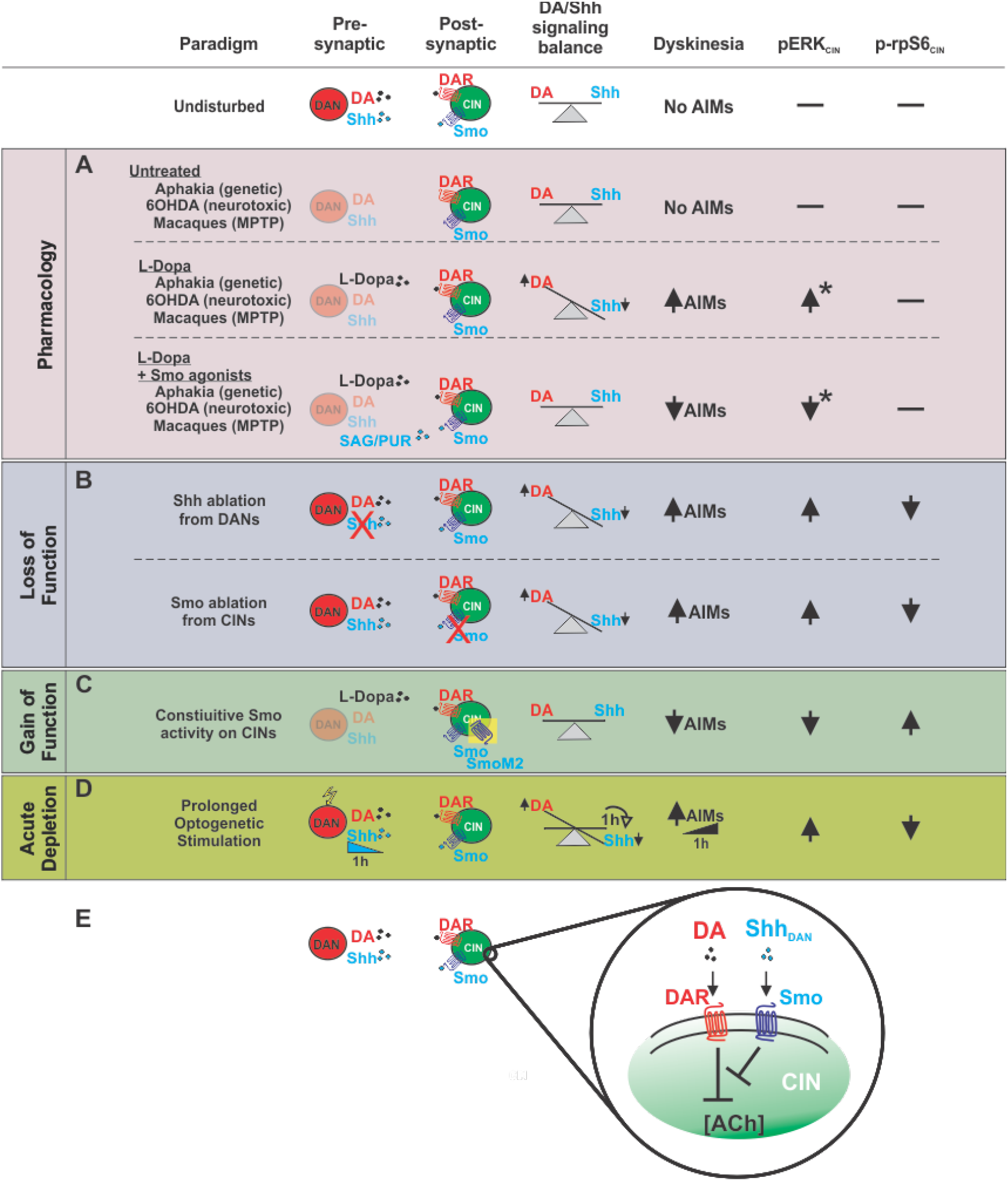
Manipulation of Shh signaling between DANs and CINs modulates LIDs, pERK, and p-rpS6 across multiple paradigms spanning timescales and degrees of DAN denervation. **(A-D)** Across the animal models of PD and experimental paradigms utilized in this study, a relative imbalance between Shh and DA signaling onto CINs was consistently observed to induce LID-like pathology. This pathology was accompanied by decreases in neuronal activity marker p-rpS6 and increases in LID histological marker pERK among CINs of the DL striatum. States defined by relatively low Shh and high DA signaling onto CINs facilitated LID. Postsynaptic manipulations of Smo on CINs produced effects mimicking the complementary pharmacological and pre-synaptic manipulations suggesting CINs are the relevant cell type through which Shh-dependent modulatation of LIDs occurs. DANs are the physiological source of Shh on which CINs depend and ablation of Shh from DANs produced LID-like behavior and changes in CIN pERK and p-rpS6 levels. Further, not only was acute Smo pharmacology sufficient to rapidly attenuate LID following SAG administration, but prolonged optogenetic stimulation of DANs led to a diminishment of Shh signaling within the course of a single hour and produced behavior resembling that observed in LID. The results obtained via this novel LID-inducing paradigm suggest that the relative balance between DA and Shh signaling onto CINs is not only perturbed following drastic denervation of DANs, but can vary at shorter timescales. **(E)**. The congruency of the findings derived from these distinct paradigms indicates that the Shh/Smo signaling pathway counteracts DA signaling in CIN. *Changes in pErk determined from examinataion of murine models.

By identifying the commonality among these diverse paradigms which differ greatly in their etiology and in the extent to which they model PD pathophysiology, we were able to pinpoint that the relative balance between Shh and DA signaling onto CINs is a critical factor in LID. Thus, our data indicate that signaling from Shh_DAN_ to SmoCIN prevents LID formation and expression through a mechanism that impinges on CIN physiology, and thus acts upstream of previously reported LID-attenuating mechanisms such as muscarinic-dependent LTD of D1 MSNs [23] or possible nicotinic receptor desensitization [22].

Our data reveals that reduction of Shh_DAN_ is a key mechanism by which DAN degeneration results in abberant cholinergic physiology. These findings concord well with the recent reappraisal of Barbeau’s DA and ACh “seesaw” [60, 67] and potentially reconcile the perplexing observation that ablation and inhibition of CINs [17, 22] as well as augmentation of cholinergic signaling can attenuate LID [22, 23]. It is well understood that DAN degeneration and the resulting hypodopaminergic state induces aberrant CIN function which in turn is a key determinant of LID induction [19, 20, 68]. However, in contrast to the long held assumption that DAN degeneration results in overall increased cholinergic (ACh) tone [69], it was recently observed that animals which underwent DAN ablation show reduced ACh tone compared to controls [60]. The pre-existing supposition that ACh is elevated following DAN degeneration can in part be explained by the fact DAN lesioned animals showed a greater reduction in DA as compared to Ach. Thus DAN degeneration appears to produce a hypercholinergic state relative to simultaneous DA signaling following DAN denervation, but a hypodopaminergic/hypocholinergic state compared to the healthy basal ganglia. This net reduction in ACh following DAN loss is in line with our previous observation that Shh_DAN_^-/-^ animals exhibit an eightfold reduction in striatal ACh concentration compared to controls [38]. Concordant, here we find that CINs of the DLS in Shh_DAN_^-/-^ animals show decreased expression of neuronal activity marker p-rpS6 which can be normalized by agonists of the Shh effector Smo or SmoM2 expression on CINs. Since ablation of CINs in the *AK*^-/-^ model has been shown to block LID [17] and forced activation of CINs by optogenetic [22] or chemogenic [12] means can induce LID, it appears the unphysiological CIN activity which emerges in LID following dopaminergic denervation bestows a LID-inductive function onto CINs which can be disengaged through their ablation. These previous observations together with our current findings that the pharmacological activation of Smo or conditional SmoM2 expression in CINs of the hypodopaminergic brain of *AK*^-/-^ mice can block LID, suggests that the interruption of Shh_DAN_ to Smo_CIN_ signaling is not merely permissive for LID but is a critical mechanism by which LID-inducing CIN pathology is set in motion following the loss of DANs.

The evidence that Shh/Smo signaling acts across multiple time scales via both rapid G-protein-dependent signaling and slower transcriptional regulation of target genes [58, 70] suggests that Shh_DAN_ to SmoCIN signaling might impinge on several homeostatic mechanisms which help maintain physiological CIN activity and thus become dysregulated following DAN degeneration and the loss of Shh_DAN_. For example, 6-OHDA lesioning or diphtheria toxin mediated ablation of DANs results in decreased spontenous activity and excitability of CINs despite a simultaneous reduction of inhibitory D2 receptor (D2R) activity [19, 29, 60]. In these models Shh_DAN_ levels must be depressed due to DAN degeneration. Consistent, we found reduced expression of the CIN activity marker p-rpS6 in the DLS of 6-OHDA lesioned mice, which could be restored to normal levels by administration of Smo agonist. Consistent with the current findings, reduced Smo signaling is also known to activate transcription of the muscarinic autoreceptor M2 [38]. Therefore, changes in transcriptional regulation of muscarinic autoreceptor expression following a loss of Shh_DAN_ to SmoCIN signaling could underlie the long term depression of tonic CIN activity previously observed in 6-OHDA or diphtheria toxin lesion studies [19, 60].

We additionally found that acute Smo agonist exposure can reduce pERK and boost p-rpS6 levels in CINs within minutes through a mechanism that requires Smo expression on CINs. This observation suggests that SmoCIN might also regulate CIN physiology through more rapid canonical G-Protein Coupled Receptor signaling. Smo signaling appears to functionally oppose the well established D2R mediated inhibition of CINs [29]. This functional opposition between DA and Shh signaling on CINs was also evident at the level of LID expression in the pharmacological and genetic paradigms of the current study: Consistent across paradigms, increasing Smo activity relative to DA signaling in CINs attenuated LID while increasing DA signaling by L-Dopa or DAN stimulation facilitated LID (Fig. 6 A-D). Concordant, we found that the dose of Smo agonist needed to counteract LID was dependent on the dose of L-Dopa used to induce LID, further suggesting that these pathways inhibit each other in a graded and interdependent manner in CINs (extended data Fig. 2).

In a recent study, 6-OHDA lesion and subsequent L-Dopa administration produced an increase of tonic CIN firing beyond that observed at normal levels [19]. This observation suggests that DAN degeneration results in a rearrangement of DA effector mechanisms in CIN from mainly inhibition via D2R - to predominantely activation via likely D5R – engangement [29, 34]. This L-Dopa dependent activation of CIN in the hypodopaminergic brain was determined to arise from reduced calcium activated potassium (SK) channel activity among CINs which could not be restored by L-Dopa or modulators of glutamatergic or GABAergic signaling [19]. The pharmacological normalization of SK channel activity attenuated LID, presumably by restoring CIN properties disrupted through DAN degeneration [19]. Interestingly, SK channels are activated by calcium entry and are inhibited by muscarinic M2/M4 autoreceptor signaling [26, 71, 72], which is increased in the absence of Shh_DAN_ to SmoCIN signaling [38]. Thus, reduced SmoCIN activity following the loss of Shh_DAN_ could depress SK channel activity and cause, in part, the altered electrophysiology observed in CINs in response to DAN degeneration [19].

Together, our data demonstrates that Shh_DAN_ to SmoCIN signaling attenuates LID in the L-Dopa treated parkinsonian brain and thus provides a strong rationale for exploiting effectors of Smo signaling in CINs for the development of anti-LID medication. Further, it will be of interest to explore whether other side effects of L-Dopa treatment, like impulse control diseases, or L-Dopa treatment refractory PD signs like cognitive deficits [73, 74] can be attenuated by manipulating the balance between Shh_DAN_/Smo_CIN_- and DA-signaling. Finally, the continued exploration of how Shh_DAN_ influences physiological CIN processing, particularly through acute temporal fluctuations in Smo signaling, is of interest for studies of basal ganglia dependent learning and motor coordination.

## Acknowledgements

This work was supported by the American Parkinson Disease Association, Research Foundation of the City University of New York, and NIH NS095253 to A.H.K., and in part by NIH R25GM56833 (P.I. Mark Steinberg) via a L.M.’s “RISE” fellowship. L.M., D.R.Z, S. U-C., and L.S. acknowledge the administrative, travel, and mentoring support they received through the biology graduate programs of the Graduate Center of the City University of New York and the CUNY School of Medicine. A.H.K. thanks John Martin for active mentoring and support. We thank Luis Gonzalez-Reyes for help with statistical analyses. We are grateful to Y.J.Z., L.H., C.Y.L., and X.R.L for the care of the non-human primates. We thank Paul Forlano, Christoph Kellendonk, John Martin, Luis Reyes-Gonzalez and Stephen Rayport for their critical comments on the manuscript.

## Contributions

L.M., D.R.Z. and A.H.K designed all the mouse experiments. A.H.K. wrote the paper. L.M. and D.R.Z. performed all mouse experiments and surgeries with contributions from S.U-C., L.S.. and H.R. in data analysis. H.R. and E.F. analyzed the neurotoxic LID paradigm. D.R.Z. developed the optogenetic Shh exhaustion paradigm. C.Q., L.Q. and E.B. conducted the NHP experiments and analyzed the associated data. E.B. contributed to manuscript writing. A.H.K. supervised all aspects of the study.

## Corresponding author

Andreas H. Kottmann.

## Ethics declarations

The authors declare no competing interests.

## Data availability

All data for this study are available from the corresponding author upon request.

---

**Extended Data Fig. 1:**
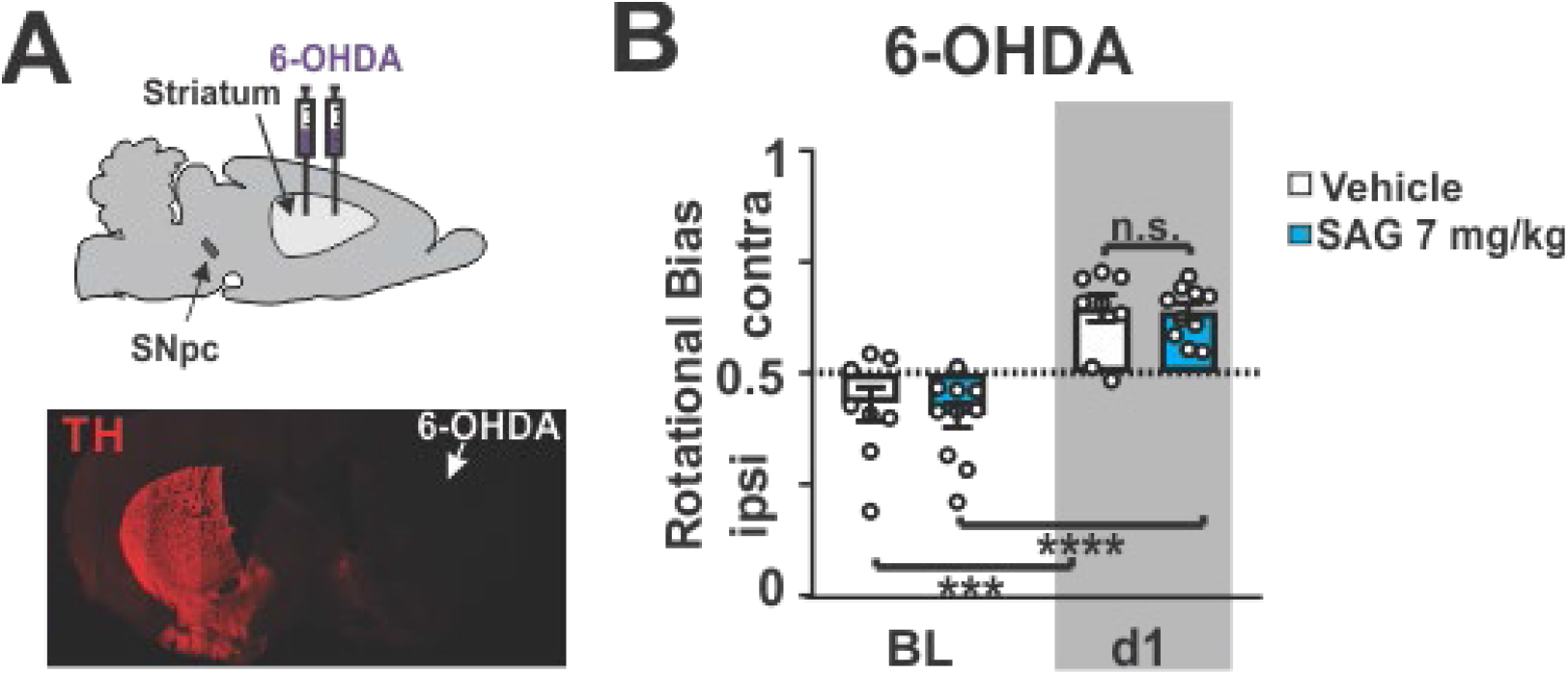
Histological and functional validation of the unilateral 6-OHDA lesion model. **(A)** 6-OHDA was injected unilaterally into the striatum at two positions along the anterior-posterior axis. Immunohistochemical staining of tyrosine hydroxylase (TH) following 6-OHDA injection revealed an almost complete loss of dopaminergic projections in the lesioned hemisphere on coronal sections. **(B)** Quantification of rotational bias as a ratio of contra-to ipsilateral turns. Dotted line signifies the absence of turning bias. Mice with 6-OHDA lesions (baseline: BL, white bar) turned ipsilateral to the lesion, indicative of hypodopaminergia in the lesioned hemisphere. Upon L-Dopa (5 mg/kg) dosing, mice turned contralateral towards the lesion, suggesting dopamine hypersensitivity had formed in the lesioned hemisphere (d1, white bar). Co-injection of L-Dopa with Smo agonist SAG (7 mg/kg, blue bars) did not alter turning bias caused by L-Dopa alone. (n = 8–9; RM two-way ANOVA time effect: F _(2,50)_ = 20.19, P<0.0001. Post hoc Bonferroni’s test: *** P<0.001 BL vs. day 1).

**Extended Data Fig. 2:**
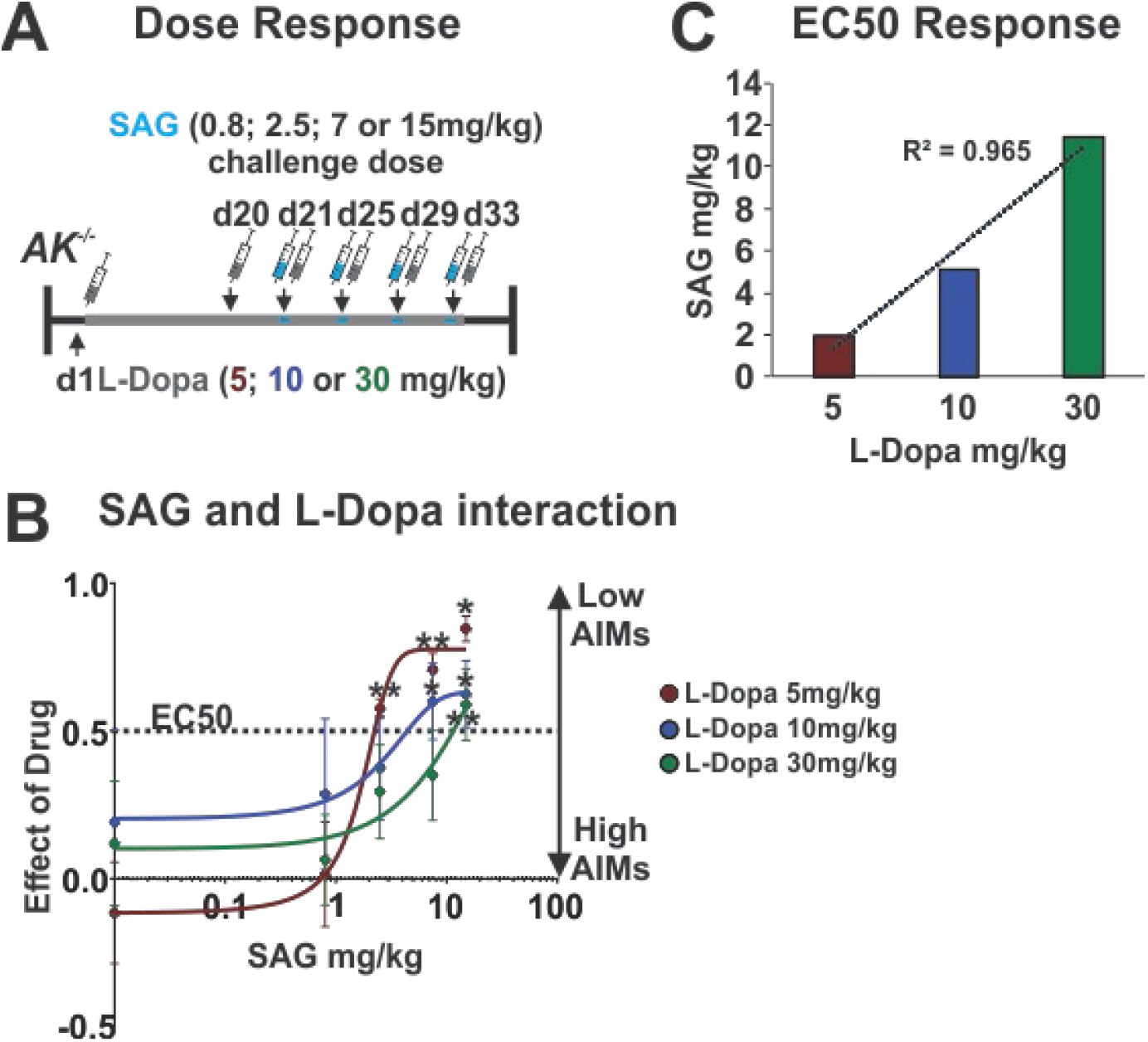
SAG dose needed to attenuate LID is positively and linearly correlated with L-Dopa dose. **(A)** LID were induced in 3 groups of *AK*^-/-^ mice injected daily for 20 days with either either 5, 10 or 30 mg/kg L-Dopa. In probe trials at days 21, 25, 29, and 33, the dose-dependent degree of LID attenuation by SAG was tested using a within-subject escalation strategy including three-day SAG washouts (L-Dopa only) inbetween probe trials. SAG was serially administered across the probe trials at 4 different concentrations (0.8; 2.5; 7.5; and 15 mg/kg) in each L-Dopa dose group (5, 10 or 30 mg/kg) **(B)** In each L-Dopa concentration group, the attenuation of LID by SAG was SAG dose-dependent. From this data we estimated the SAG concentration that resulted in half-maximal (EC50) inhibition of LID across the L-Dopa groups. Linear regression lines are plotted to best fit the data points. Stars represent statistically significant difference from baseline AIMs when L-Dopa and vehicle were administered. (n = 8 for each drug condition; unpaired two-tailed Student’s t test * P<0.05, ** P<0.01. n.s. indicates P>0.05. Graph is plotted as mean +/- SEM.) **(C)** Analysis revealed a positive correlation between the dose of SAG needed to attenuate AIMs and the concentration of L-Dopa used to induce LID (R^2^ = 0.965). These results suggest that the degree of LID severity correlates positively with the degree of Shh/Smo- and DA-signaling imbalance.

**Extended Data Fig. 3:**
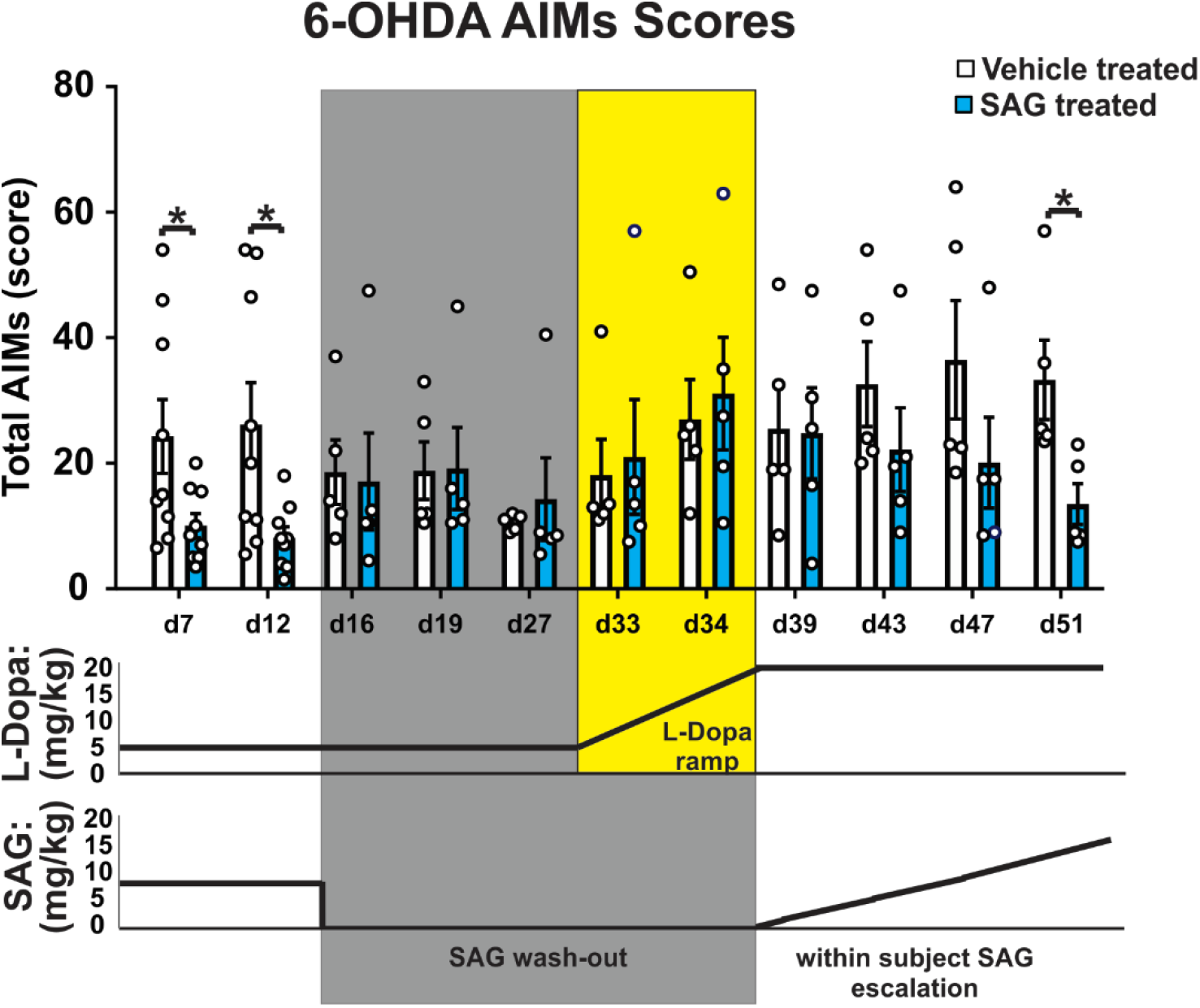
AIMs can be repeatedly attenuated or reinstated, respectively, through sequential dosing of L-Dopa with or without SAG. Daily co-injection of L-Dopa and Smo agonist SAG (5 mg/kg) for seven days significantly attenuated LID compared to L-Dopa only injected controls in 6-OHDA animals. Additional days of L-Dopa and SAG co-administration did not further reduce LID (d12). Terminating SAG dosing on day 16 while continuing L-Dopa treatment resulted in an immediate reappearance of LID to levels seen in controls. Severity of reinstated LID remained sensitive to L-Dopa concentration such that a gradual increase in L-Dopa dosing led to greater AIM scores with no difference in kinetics or absolute intensity compared to controls (d33–d34). Reinstated LID could be attenuated again in a dose-dependent manner through within-subject escalation of SAG on days 39–51. During this within-subject SAG escalation, three-day SAG washout periods during which only L-Dopa was administered were included between days of increasing SAG dose administration (n = 9 for d 7, 12; n = 5 for d 16–51; Paired two-tailed student’s t test: * P<0.05 treatment vs. vehicle. n.s. indicates P > 0.05). Bar graph is plotted as mean +/- SEM.

**Extended data Fig. 4:**
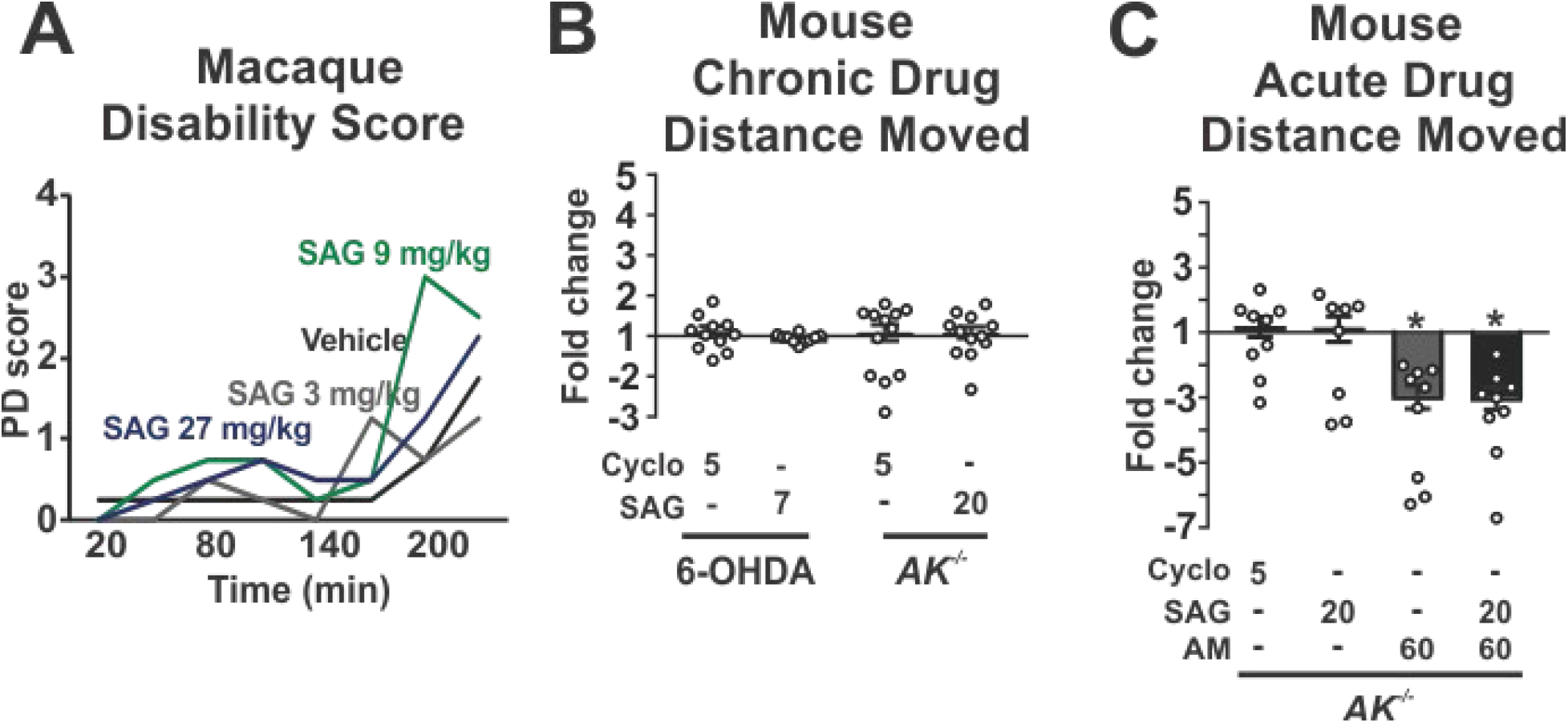
SAG treatment does not curtail the anti-parkinsonian benefit of L-Dopa. **(A)** Parkinsonian disability score of Parkinsonian Macaques across a four-hour time course after receiving L-Dopa together with either vehicle or SAG (3, 9, and 27 mg/kg; n = 4 per condition). Scoring involved evaluating a range of movements including bradykinesia, postural abnormality, and tremor, yielding a maximum global parkinsonian disability score of 10 (Methods). There was no effect of SAG on the Parkinsonian disability score signifying that SAG did not diminish the akinetic benefit of L-dopa treatment. Error bars are excluded for clarity. **(B)** Fold change of distance moved in an “Open Field” arena following daily co-injection of L-Dopa and either Cyclo (5 mg/kg) or SAG (7 mg/kg for 6-OHDA mice or 20 mg/kg in *AK*^-/-^ mice) in 6-OHDA (day 14 of treatment; n = 8) and *AK*^-/-^ (day 26 of treatment; n = 12) mice. Results are reported as fold change over vehicle-treated controls. Cyclopamine or SAG co-injection with L-Dopa in 6-OHDA treated or *AK*^-/-^ mice did not alter locomotion activity compared to L-Dopa + vehicle controls **(C)** Fold change of distance moved on day 20 of daily L-Dopa injections *AK*^-/-^ mice that received an acute dose of either Cyclo (5 mg/kg; n = 9), SAG (20 mg/kg; n = 8), Amantadine (AM, 60 mg/kg; n=9), or a combination of AM (60 mg/kg) and SAG (20 mg/kg; n = 8). AM significantly reduced the anti-akinetic benefit of L-Dopa but SAG did not (unpaired two-tailed student’s t test: * P<0.05 for treatment vs. vehicle. n.s. indicates P>0.05). All bar graphs are plotted as mean +/- SEM.

**Extended Data Fig. 5:**
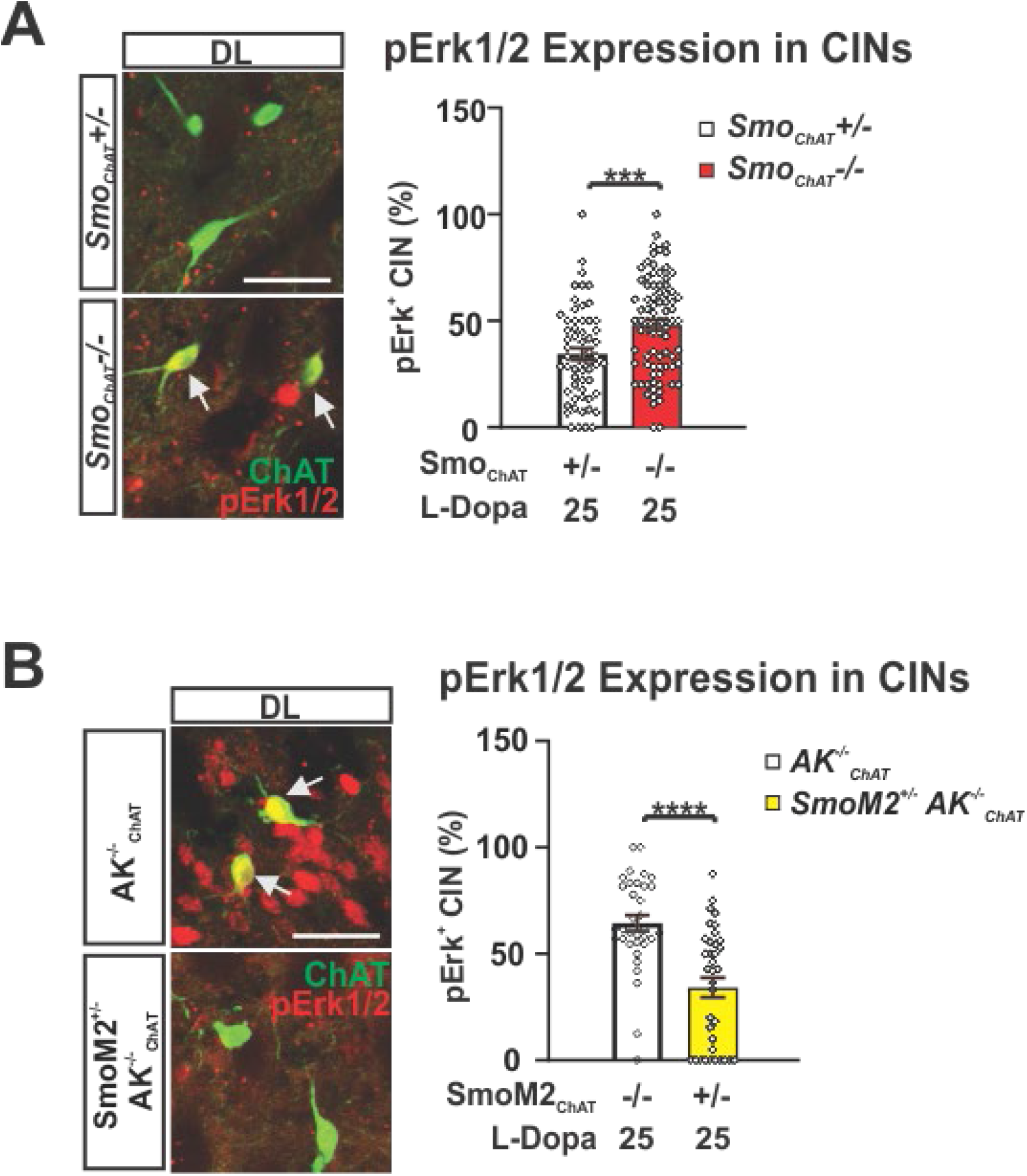
Conditional Smo ablation increases prevalence of pErk^1/2^-positive CIN while conditional SmoM2 expression decreases prevalence of pErk^1/2^-positive CINs in the DLS. **(A)** Representative images of pErk^1/2^ (red) co-localization (arrow) with CINs (ChAT, green). Quantification and conmparison of pErk prevalence in the DLS of *Smo_ChAT-cre_*^-/-^ and *Smo_ChAT-cre_^-/+^* heterozygous control littermates was performed following daily dosing with L-Dopa (25 mg/kg) for 20 days (n = 6–7 per genotype, 35–38 CINs each). Unpaired two-tailed student’s t test *** P<0.001 control vs. mutant. **(B)** Representative images of pErk^1/2^ (red) co-localization (arrows) with CINs (ChAT, green). Quantification and comparison of pErk prevalence in the DLS of *SmoM2*^+/-^ *AK*^-/-^_*ChAT-Cre*_ and *AK*^-/-^_*ChAT-Cre*_ littermate controls was performed following daily dosing with L-Dopa (25 mg/kg) for 20 days (n = 8–12, 35–38 CINs each; unpaired two-tailed student’s t test **** P<0.0001 control vs. mutant. All bar graphs are plotted as mean +/- SEM. Scale bar = 50 μm.

**Extended Data Fig. 6:**
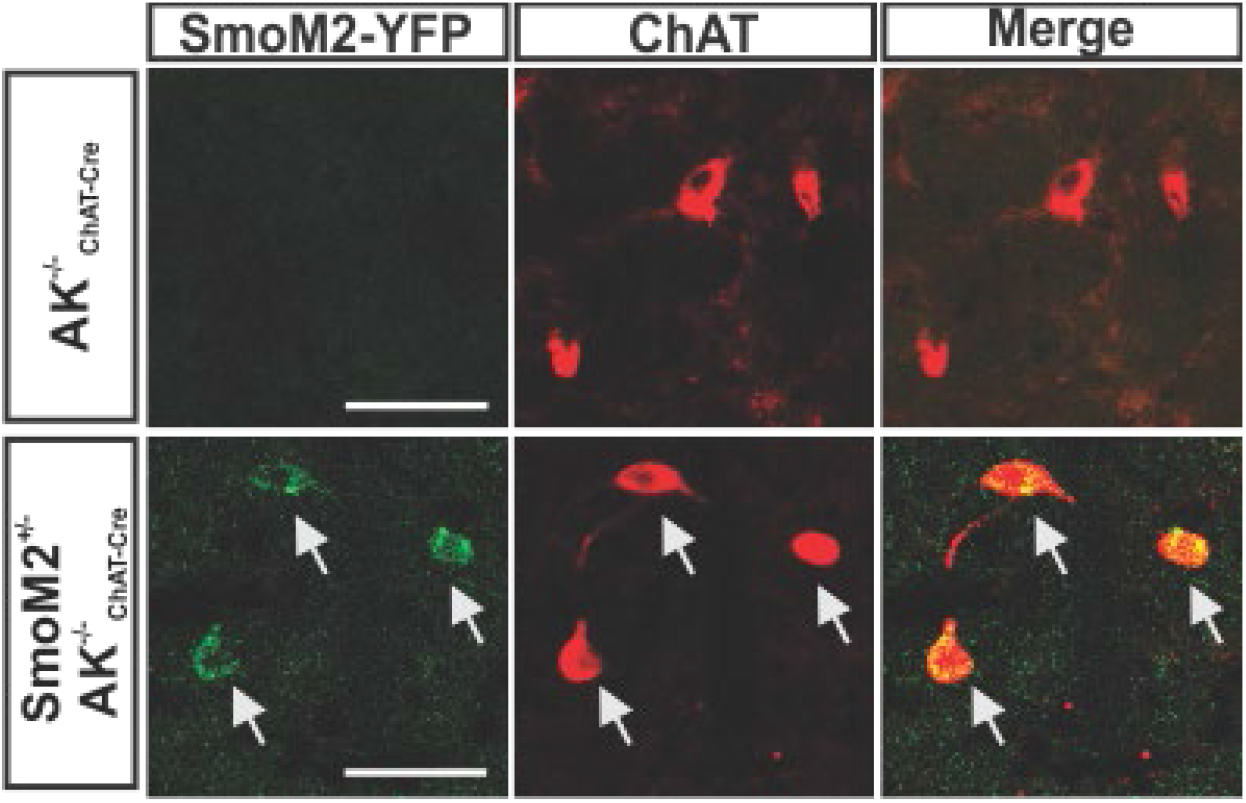
Conditional expression of SmoM2 in CINs of *AK*^-/-^ mice. Images of the DLS showing eYFP-tagged SmoM2 expression selectively in CINs of *SmoM2*^+/-^ *AK*^-/-^_*ChAt-Cre*_ mice (arrows) but not in *AK*^-/-^_*ChAT-Cre*_ littermate controls. CINs identified through co-labeling with ChAT. Scale bar = 50 μm.

**Extended Data Figure 7:**
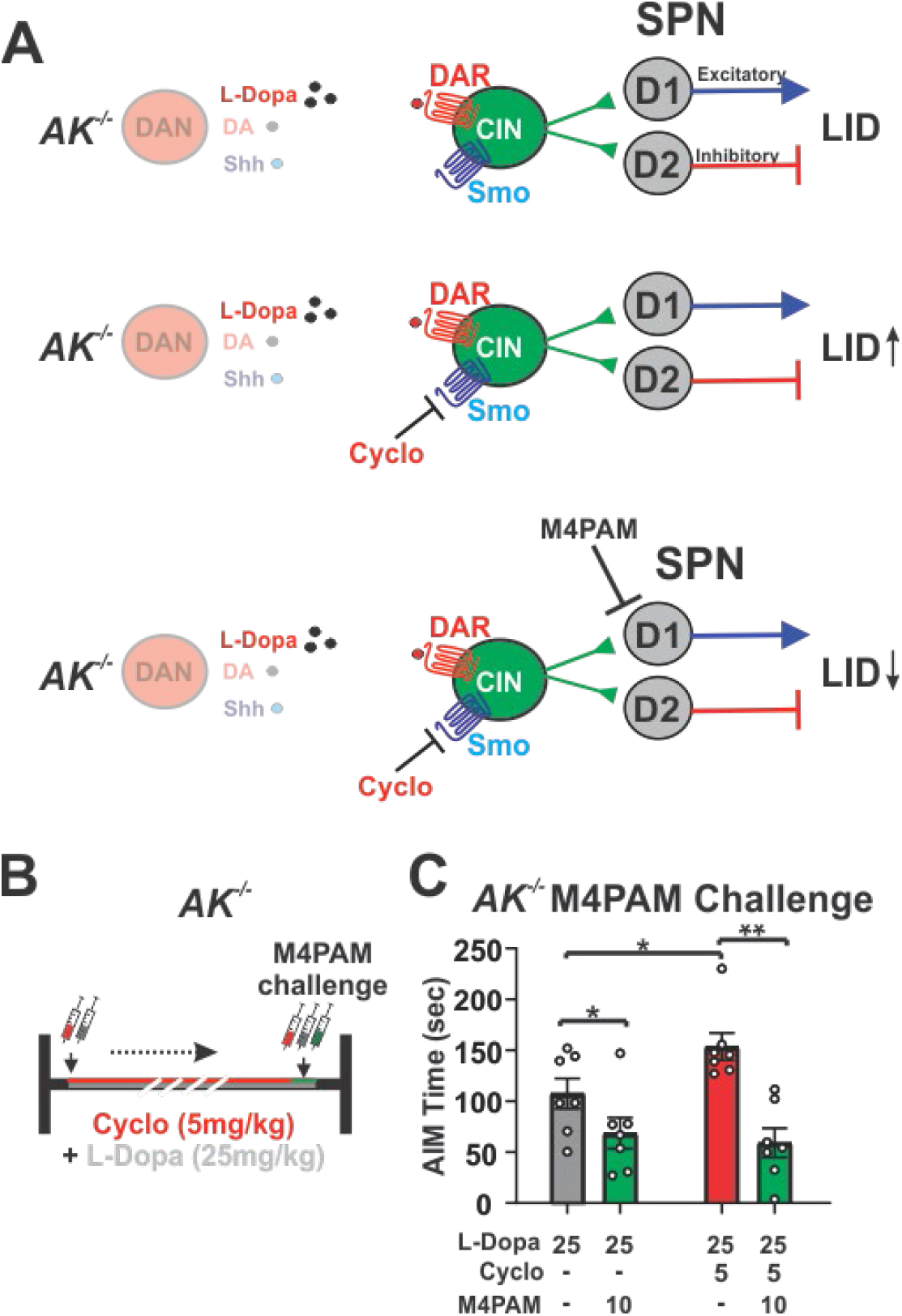
Shh_DAN_ to Smo_CIN_ signaling attenuates AIMs via a mechanism upstream of cholinergic influence on striatal output pathways. **(A**) Muscarinic acetylcholine receptor 4 positive allosteric modulators (M4PAMs) attenuate AIMs by boosting cholinergic signaling onto striatal D1 medium spiny projection neurons (D1-MSNs) and facilitating their long-term depression (LTD) [23]. The process of D1-MSN LTD reduces excitatory motor output from the basal ganglia and thus leads to a net reduction of LID. This observation concords well with our finding that increased p-rpS6^**240/244**^ levels in CINs correlate with reduced AIMs (Fig. 4). The characterized inhibition of LID by M4PAMs allowed us to test whether Smo-mediated modulation of AIMs in *AK*^-/-^ mice occurs upstream or downstream of cholinergic action on D1-MSNs. **(B)** Dosing schedule of *AK*^-/-^ mice with L-Dopa (25 mg/kg) and vehicle or Cyclopamine (5 mg/kg) for 20 days. Mice were challenged with a single dose of M4PAM VU0467154 (10mg/kg) alone or in combination with Cyclopamine (5mg/kg) on the final day. **(C)** We found that M4PAMs can attenuate Cyclopamine-caused facilitation of AIMs (red bar) in *AK*^-/-^ animals down to the same levels achieved by M4PAMs in *AK*^-/-^ mice treated with L-Dopa alone (grey bars). This finding indicates that Smo-mediated modulation of AIMs occurs upstream of M4PAM mediated modulation of AIMs (n = 7; paired two-tailed Student’s t test P<0.05, * P<0.05, ** P<0.01 veh vs. M4PAM; Unpaired two-tailed Student’s t test P<0.05, * P<0.05 vehicle vs. Cyclopamine). Bar graph is plotted as mean +/- SEM.

**Extended Data Figure 8:**
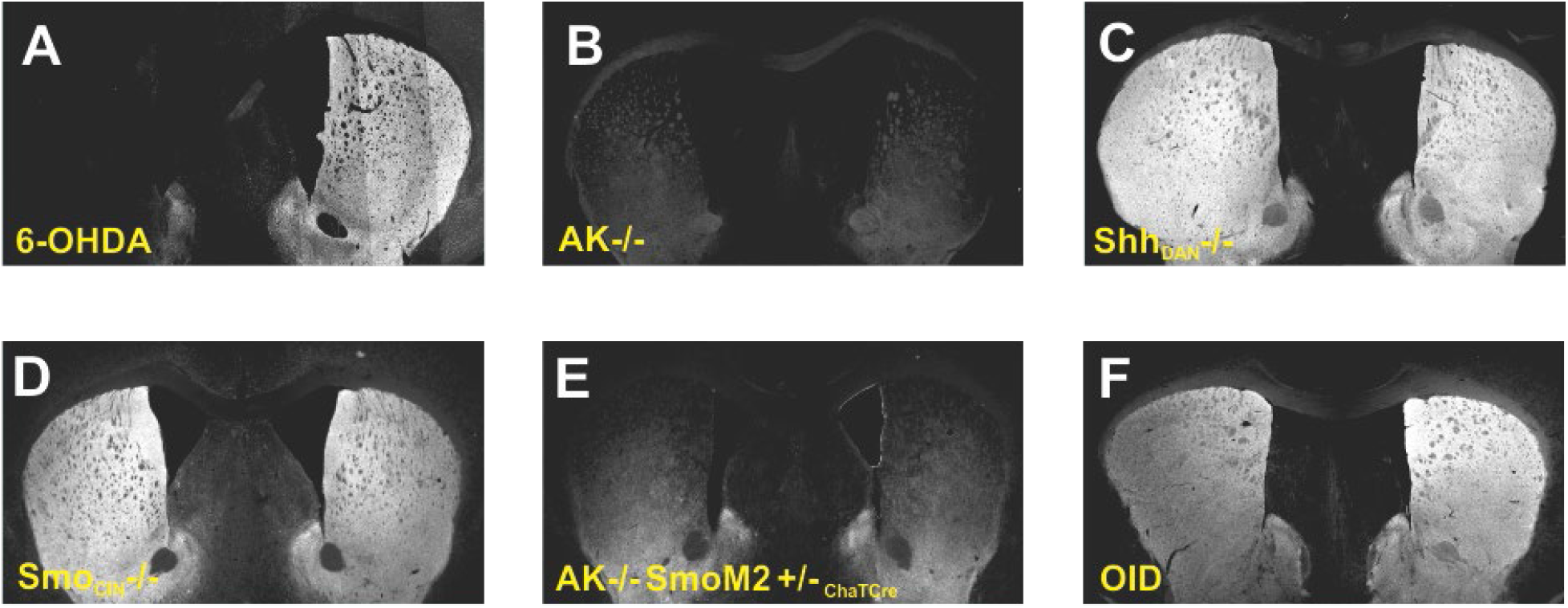
TH fiber density does not predict LID formation and expression. Tyrosine hydroxylase (TH) fiber density was examined as a proxy for the integrity of dopaminergic projections across the mouse models utilized in this study. **(A, B)** As reported previously, 6-OHDA injection and the ablation of Pitx3 in *AK*^-/-^ animals resulted in a severe reduction of dopaminergic projections to the striatum ([48, 76], suggesting that all co-transmitter signaling from DANs was reduced in these projection areas. **(C, D)** In contrast, there was no clear diminishment of TH fiber density in the striatum of *Smo_ChAT-Cre_*^-/-^ or *Shh_DAN_*^-/-^ animals. **(E)** Conversely, crossing the *SmoM2_ChAT-Cre_*^+/-^ allele into *AK*^-/-^ animals did not restore the severely diminished TH fiber density observed in the DLS of *AK*^-/-^ animals. **(F)** Prolonged burst activity forced by optogenetic stimulation (one hour per day for 10 days) did not reduce TH fiber density. Together, these observations suggest that overt dopaminergic denervation might not be necessary to allow LID formation or expression and emphasize the importance of Shh_DAN_ to SmoCIN signaling among all dopaminergic signaling modalities for AIMs suppression.

**Extended Data Fig. 9:**
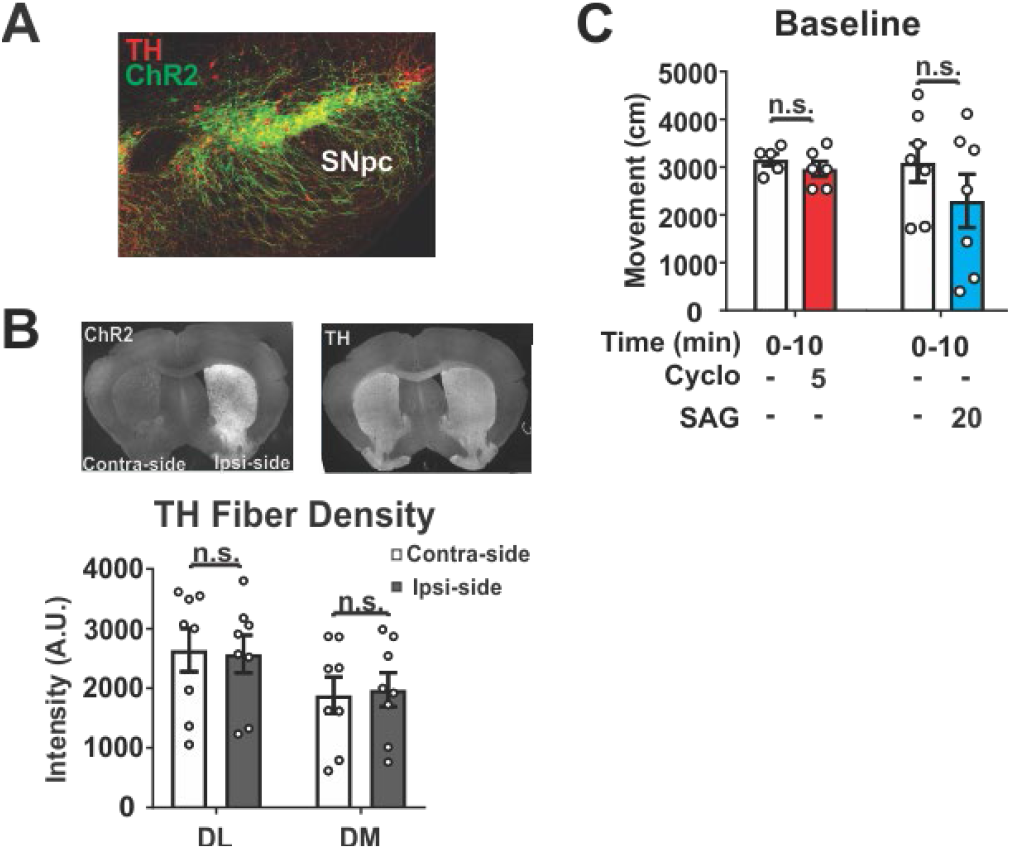
Daily repeated, long-term optogenetic stimulation of DANs does not reduce dopaminergic fiber density in the striatum. **(A)** Co-localization of the channelrhodopsin (ChR2):: eYFP (green), fusion protein and TH (red) in the SNpc of Dat-Cre^+/-^ animals. **(B)** Representative images and quantification of TH fiber density after daily, hour long optical burst stimulation of DANs for 10 days. Comparison is made between the hemisphere ipsilateral to ChR2 expression and the contralateral striatum (n = 8; n.s. indicates P > 0.05).

## Methods

### Mice

All mice were maintained on a 12 h light/dark cycle, with ad libitum food and water. Animal use and procedures were in accordance with the National Institutes of Health guidelines and approved by the Institutional Animal Care and Use Committees (IACUC) of the City University of New York. All experiments were carried out in young adult animals beginning at 2 months of age and weighing between 22 – 28 g unless stated otherwise. Males and females from a mixed CD1 and C57/BL6 background were used for all experiments in approximately equal proportions.

*C57/Bl6* mice, JAX # 000664, were purchased from Jackson Laboratory.

*AK*^-/-^ mice are homozygous *Pitx3^ak/ak^* mice and were generously provided by Dr. Un Jung Kang (NYU), propagated by homozygous breeding and genotyped as previously described in [1, 2]).

*Shh_DAN_*^-/-^ mice and controls are produced by crossing Shh-nLacZ^L/L^ females with Shh-nLacZ^L/+^/DatCre males resulting in Shh-nLacZ^L/L^/Dat-Cre experimental (*Shh_DAN_*^-/-^) - and Shh-nLacZ^L/+^/Dat-Cre (*Shh_DAN_*^-/+^) control mice as previously described in [3].

*Smo_ChAT-Cre_*^-/-^ mice and controls are produced by crossing Smo^L/L^ females with Smo^L/+^ /ChAT-IRES-Cre males resulting in Smo^L/L^ /ChAT-IRES-Cre (*Smo_CIN_*^-/-^) experimental – and Smo^L/+^/ChAT-IRES-Cre (*Smo_CIN_^-/+^*) control mice. Smo ^L/+^ (JAX # 004526; [4]) and ChAT-IRES-Cre (JAX # 006410; [5] mice were purchased from Jackson Laboratory.

*SmoM2^+/-^_ChAT-Cre_ AK^-/-^* mice are ^L^STOP^L^SmoM2-YFP/ChAT-IRES-Cre /Pitx3^ak/ak^ mice and are maintained by crossing Pitx3^ak/ak^ females with triple heterozygous ^L^STOP^L^SmoM2-YFP/ChAT-IRES-Cre /Pitx3^ak/+^ males resulting in ^L^STOP^L^SmoM2-YFP/ChAT-IRES-Cre /Pitx3^ak/ak^ (*SmoM2^+/-^_ChAT-cre_ AK*^-/-^) experimental – and ChAT-IRES-Cre /Pitx3^ak/ak^ (*_ChAT-Cre_ AK*^-/-^) – control mice. Tissue specific activation of SmoM2 was confirmed by eYFP expression. ^L^STOP^L^SmoM2-YFP mice (JAX # 005130; [6]) were purchased from Jackson Laboratory.

### Genotyping

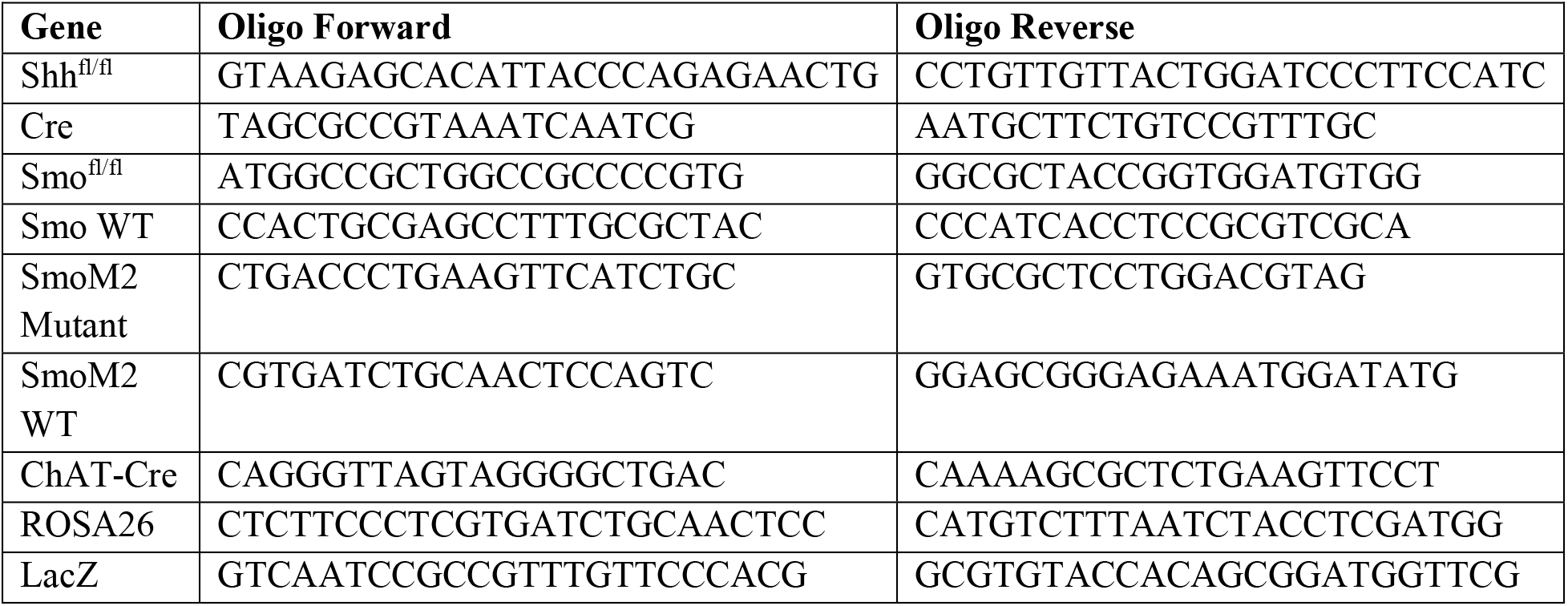

### Macaques

The non-human primate experiments were performed in accordance with the European Union directive of September 22, 2010 (2010/63/EU) on the protection of animals used for scientific purposes in an AAALAC-accredited facility following acceptance of study design by the Institute of Lab Animal Science (Chinese Academy of Science, Beijing, China). The four Macaca fascicularis male monkeys (Xierxin, Beijing, PR of China) were housed in individual cages allowing visual contact and interactions with other monkeys in adjacent cages. Food and water were available *ad libitum*. Animal care was supervised daily by veterinarians skilled in the healthcare and maintenance of NHPs.

### Antibodies

Anti-tyrosine hydroxylase (1:500; RRID: AB_657012) and anti-ChAT (1:100; RRID: AB_144P) were purchased from Milipore. Anti-phospho-Erk1/2 (1:400; RRID: AB_9101), and anti-phospho-rpS6 (ser240/244; RRID: AB_5364) were from Cell Signaling Technology. Anti-NeuN (1:200; RRID: AB_MAB377) from Chemicon International. Various Alexa Fluor antibodies were used at 1:250 dilutions for immunohistochemistry (Jackson Immuno Research).

### Drugs

In mice, all pharmacological agents were administered via intraperitoneal (i.p.) injection. Animals were treated daily in a volume of 10 mL/kg of body weight with a combination of L-Dopa (5-25 mg/kg; Sigma-Aldrich D1507) and the peripheral L-amino acid decarboxylase antagonist, benserazide (12.5-20 mg/kg; Sigma-Aldrich B7283) diluted in 0.9% sterile saline (referred to as just “L-Dopa” and specified doses described in main text). Sonic Hedgehog agonist (SAG Carbosynth Limited FS27779; 0.8-20 mg/kg) and antagonist (Cyclopamine Carbosynth Limited FC20718; 2.5-5 mg/kg) were dissolved in DMSO and brought into solution with 45% HPCD (Sigma Aldrich H107) in 0.9% sterile saline for 6-OHDA and *AK*^-/-^ AIM studies. SAG-HCl (Carbosynth Limited FS76762) was diluted in 0.9% sterile saline and used for the *Shh_DAN_^-/-^* and *Smo_CIN_*^-/-^ mouse studies. Both Cyclopamine and SAG were administered 15 minutes before L-Dopa injections. Amantadine hydrochloride (Sigma Aldrich A1260) was dissolved in 0.9% sterile saline and given 100 minutes prior to L-Dopa administration. M4PAM (VU0467154 StressMarq Biosciences SIH184) was dissolved in 0.9% sterile saline and given at the same time as L-Dopa. Purmorphamine (ABCam AB120933) was dissolved in a cocktail of Polyethylene glycol (PEG) and ethanol in PBS and given 15 minutes before L-Dopa dosing. Control mice were treated with carrier (DMSO, HPCD etc.) in similar proportions to the treatment drugs.

### Mouse unilateral 6-OHDA model [9]

Mice were anesthetized with a mixture of ketamine (80 mg/kg) and xylazine (12 mg/kg) administered via i.p. and the surgical field was prepared with betadine. Bupivacaine (Marcaine Sigma Aldrich B5274), a local anesthetic, was subcutaneously (s.c.) injected near the incision site. Bregma was visualized with 3% hydrogen peroxide. Two unilateral injections (2 x 2 μl each) of 6-OHDA were administered into the left striatum using the following coordinates based on the Mouse Brain Atlas by Paxinos and Franklin (2001): Anteroposterior (AP) +1.0 mm; Lateral (ML), + 2.1 mm; Dorsoventral (DV), −2.9 mm; and AP +0.3 mm; L +2.4 mm; DV −2.9 mm. 6-OHDA-HCl (Sigma Aldrich H4381; 3.0 mg/ml) was dissolved in a solution containing 0.2 g/L ascorbic acid (Sigma Aldrich A92902) and 9 g/L NaCl and injected via a Hamilton syringe with a 33-gauge needle attached to a micro-syringe pump (World Precision Instruments) at a rate of 0.4 μl/min and left in place for 3 minutes afterwards before being retracted slowly. Following surgery, animals were injected with 5% sucrose (10 ml/kg, s.c.) and saline (10 ml/kg, i.p.) and recovered on a heating pad. To avoid dehydration and weight loss, hydrogel pouches and hi-fat chow were given to the mice ad lib. Behavioral testing and drug treatment began three weeks following surgery. Lesion assessments were quantified by ipsilateral paw-use bias in the cylinder test and verified histologically, post-mortem, by quantification of tyrosine hydroxylase (TH) fiber density at the end of experiments. Only animals with 70% or more of TH positive fiber depletion were included in the analyses.

### Forelimb cylinder test

The cylinder test was used to assess the anti-akinetic effects of L-Dopa in 6-OHDA lesioned mice by measuring forelimb paw placement. Mice were placed in a glass cylinder (10 cm wide x 14 cm high) 30 minutes post L-Dopa injection and analyzed for 4 minutes. Two mirrors were placed in the back to observe mice from all angles. The limb use asymmetry score was expressed as a ratio of contralateral to the lesion paw use over the total number of wall contacts with both paws as described previously in [10].

### Open field test

In the open field test (OFT), whole body movements were measured for 5 min using a Noldus Ethovision XT video tracking system. The animals head, body, and tail positions were tracked in a transparent Plexiglas 4-chamber box apparatus (50 x 50 cm for each chamber). Spontaneous locomotion was quantified as distanced traveled at baseline (no drug) and 45 minutes following drug administration. Asymmetry of locomotion was calculated as a percentage of lesion ipsilateral to contralateral turning bias.

### Abnormal involuntary movements

Abnormal involuntary movements (AIMs) were scored using the established behavioral rating scales [10–12]. Briefly, there are four types of AIMs: 1) abnormal rotational locomotion, 2) axial AIMs, showing dystonic posturing, and severe twisting of the head or neck, 3) limb movements, with rapid, jerky movements of the front or hind limbs, and 4) orofacial movements of abnormal chewing, licking, grooming, or sticking out of the tongue. The four types of AIMs were scored on a severity scale of 0-4, with 0 exhibiting no abnormality and 4 showing uninterruptable abnormal movement. AIM scores were assessed by an observer blinded to the treatment, 35 minutes post treatment injection, and 20 min post L-Dopa injection. Animals were single caged during AIM assessments. “Total AIM” scores for each animal were recorded every 20 minutes, for 1 min each, over 2 hours.

### *AK*^-/-^ forelimb and 3-paw dyskinesia

Animals were recorded in a clear plastic cylinder (16 cm in diameter and 25 cm in height) 30 minutes following L-Dopa injection. Mirrors were placed behind the cylinder to allow for ventral views of the behavior. Each trial was five minutes with a 30-second habituation period followed by behavioral scoring. The duration of abnormal paw movement was quantified for each trial and included sliding and shaking of the forelimb paws on the cylinder, and “three-paw” dyskinesia (two forelimbs and one hindlimb) and “four-paw” dyskinesia (animal balances on its tail while rearing at the glass wall) was observed in extreme cases as described in [1].

### Primate MPTP model

#### Model preparation

The 1-methyl-4-phenyl-1,2,3,6-tetrahydropyridine (MPTP) intoxication protocol, chronic L-Dopa treatment, and the clinical assessments were conducted in four male macaques (Macaca fascicularis, Xierxin, Beijing, PR of China), as previously published [13, 14]. The macaques were first rendered parkinsonian with MPTP-hydrochloride (0.2mg/kg, i.v., Sigma) dissolved in saline. Daily (at 9 am) assessment of parkinsonism in home cages for 30 min by two blinded observers was done using a validated rating scale assessing tremor, general level of activity, body posture (flexion of spine), vocalization, freezing and frequency of arm movements (for each upper limb), and rigidity. Once parkinsonism was stable, levodopa (Madopar^®^, Roche, Levodopa/carbidopa, ratio 4:1) was administered twice daily for 4-5 months at an individually-tailored dose designed to produce a full reversal of the parkinsonian condition (p.o. by gavage). Over this period, animals developed severe and reproducible dyskinesia, presenting choreic–athetoid (characterized by constant writhing and jerking motions), dystonic, and sometimes ballistic movements (large-amplitude flinging, flailing movements) as seen in long-term L-Dopa-treated PD patients. SAG (3, 9, 27mg/kg) was dissolved in 10% DMSO, 45% HPCD in 0.9% saline and administered i.v.. Within subject escalation was performed with a washout period of three days between escalating doses.

Immediately after drug administration, monkeys were transferred to an observation cage (dimensions - 1.1m x 1.5m x 1.1m) as per guidelines [15]. The total duration of observation was 240 min drug exposure. We performed a battery of behavioural observations as previously described [13]. Experts blinded to the treatment observed 10-min video recordings taken every 30 min throughout the duration of the experiment and scored the severity of the parkinsonian condition using the parkinsonian disability score. The parkinsonian disability score is a combination of four different scores: (i) the range of movement score, (ii) bradykinesia score, (iii) posture score, and (iv) tremor score. These four scores are combined using formula: (4 - range of movement) + bradykinesia + postural abnormality + tremor. We rated the severity of dyskinesia using the Dyskinesia Disability Scale [15]: 0, dyskinesia absent; 1, mild, fleeting, and rare dyskinetic postures and movements; 2, moderate, more prominent abnormal movements, but not interfering significantly with normal behaviour; 3, marked, frequent and, at times, continuous dyskinesia intruding on the normal repertoire of activity; or, 4, severe, virtually continuous dyskinetic activity replacing normal behaviour and disabling to the animal.

We presented the time course of parkinsonian disability and dyskinesia scores in 30 min time bins over the 4 hour observation period. We also presented the median of the total scores of disability, dyskinesia, chorea, and dystonia at 0-2 hours following treatment. We statistically compared the parkinsonian and dyskinesia scores between different conditions using a Friedman’s test followed by Dunn’s multiple comparison test.

### Optogenetic manipulations

#### Implant construction and implantation

Implants and patch cables were constructed and polished as previously described [16]. Mice were anesthetized with isoflourene and the surgical field was prepared with betadine. Bupivacaine was injected in the scalp prior to incision. Bregma was visualized with 3% hydrogen peroxide and a craniotomy was made over the injection site. The AAV5-EF1a-DIO-hChR2(H134R)-eYFP-WPRE virus (UNC Vector Core) was injected at AP −3.2 mm, ML +1.5 mm, DV −4.3 mm by pressure injection with a pulled glass pipette and allowed to dwell for 10 minutes. A ferrule implant was placed at AP −3.2 mm, ML +1.5 mm, DV −4.2 mm, secured with metabond, and protected with a dust cap (Thor Labs). Following surgery, mice were given 0.05 mg/kg buprenex and recovered on a heating pad. At least one month was allowed for viral expression.

#### Optogenetic stimulation paradigms

One trial of stimulation consisted of a ten-minute habituation period followed by an hour of intermittent stimulation for 5 sec in which 10 msec long pulses at 60 Hz and a laser power of 20 mW as measured at the tip of the implant were produced, followed by a 30 sec pause. Stimulation was controlled via a TTL pulse from Ethovision’s software to a Doric four channel pulse generator controlling a 710 nm Laserglow DPPS laser. All sessions were recorded and analyzed with Ethovision. Pharmacological agents were given 30 minutes prior to the beginning of the trial.

#### Optogenetic-induced dyskinesia

For optogenetic-induced dyskinesia (OID) experiments, stimulation occurred in a 5L glass cylinder and consisted of one hour of intermittent stimulation as previously described. Stimulation occurred daily for the duration of the experiments. A second camera (Casio EX-FH100) recorded behavior from the side of the cylinder during the first and last ten minutes of the trial. AIMs were assessed on limb, orofacial, and axial movements using the same scoring scale developed for the unilateral 6-OHDA lesion paradigm [11]. AIMs were scored separately during stimulation and non-stimulation periods. Pharmacological agents were given 30 minutes prior to the beginning of the trial.

### Immunohistochemistry

Animals were anesthetized with pentobarbital (10 mg/ml) 30 minutes after last drug injection and transcardially perfused with 50 ml of 4% paraformaldehyde (PFA) in 0.2M Phosphate buffer PH 7.2. The brains were extracted and fixed overnight in 4% PFA and equilibrated in 30% sterile sucrose for 48 h. Brains were mounted in OTC and cryo-sections produced at 40 μm. Slices were incubated with antibodies as described in [17]. Sections were mounted in Vectashield (Vector Laboratories Inc.).

Confocal microscopy was performed using a Zeiss LMS 810 laser-scanning confocal microscope using the same acquisition parameters for experimental and control tissue for each experiment. Regions of interest (ROI)

TH immunofluorescence intensity was quantified in the striatum with MacBiophotonics Image J, and the data represented mean gray levels above background.

The fraction of pErk^1/2^ positive CINs was expressed as a percentage of pErk^1/2^ positive CIN out of the total number of CIN in each of 375 x 375 μm ROIs in the dorsolateral and dorsomedial striatum of each coronal section analyzed. Relative position of the ROI box was kept the same between genotypes and treatment groups and along the anterior posterior axis. Quantification was performed blinded to the treatment and/or genotype of the subject in three striatal slices: anterior (AP: ~1.34 mm), middle (AP: ~0.74 mm), and posterior (AP: ~0.14 mm) per animal. Co-stained and single stained cells were manually counted using FIJI/ImageJ software (NIH). Each data point reflects the fraction of pErk^1/2^ positive CIN in one ROI.

### p-rpS6^240/244^ Analysis

The mean grey value of p-rpS6^240/244^ intensity was measured in FIJI/ImageJ software (NIH) using a freehand ROI area delineating the soma of each ChAT stained CIN in a 375 x 375 mm box in the dorsolateral striatum as described in [18]. NeuN fluorescence intensity of the same cell was used for normalization of fluorescence across treatment and genotype groups. Heat map images reveal the intensity levels of p-rpS6^240/244^ fluorescence based on a 16 pseudocolor palette. 200-300 CINs were quantified per each condition.

### Statistics

No statistical methods were used to predetermine sample size.

Multiple comparisons were analyzed using 2-way, 3-way, or repeated measures (RM) ANOVA with GraphPad Prism 8.0 software, followed by post hoc Bonferroni’s test for specific comparisons. For 2-group comparisons, 2-tailed paired or unpaired Student’s *t* test was used. Nonparametric statistics were used to verify all main effects using Dunnett’s multiple comparison test or Mann-Whitney test when appropriate.

## References

1. Fearnley, J.M. and A.J. Lees, Ageing and Parkinson’s disease: substantia nigra regional selectivity. Brain, 1991. 114 (Pt 5): p. 2283–301.

2. Kordower, J.H., et al., Disease duration and the integrity of the nigrostriatal system in Parkinson’s disease. Brain: a journal of neurology, 2013. 136(Pt 8): p. 2419–31.

3. Cotzias, G.C., P.S. Papavasiliou, and R. Gellene, Modification of Parkinsonism--chronic treatment with L-dopa. N Engl J Med, 1969. 280(7): p. 337–45.

4. Lane, E.L., L-DOPA for Parkinson’s disease-a bittersweet pill. Eur J Neurosci, 2019. 49(3): p. 384–398.

5. Ahlskog, J.E. and M.D. Muenter, Frequency of levodopa-related dyskinesias and motor fluctuations as estimated from the cumulative literature. Mov Disord, 2001. 16(3): p. 448–58.

6. Calabresi, P., et al., Levodopa-induced dyskinesias in patients with Parkinson’s disease: filling the bench-to-bedside gap. Lancet Neurol, 2010. 9(11): p. 1106–17.

7. Bastide, M.F., et al., Pathophysiology of L-dopa-induced motor and non-motor complications in Parkinson’s disease. Progress in neurobiology, 2015. 132: p. 96–168.

8. Fahn, S., The medical treatment of Parkinson disease from James Parkinson to George Cotzias. Mov Disord, 2015. 30(1): p. 4–18.

9. Cenci, M.A., Presynaptic Mechanisms of l-DOPA-Induced Dyskinesia: The Findings, the Debate, and the Therapeutic Implications. Front Neurol, 2014. 5: p. 242.

10. Picconi, B., et al., Loss of bidirectional striatal synaptic plasticity in L-DOPA-induced dyskinesia. Nat Neurosci, 2003. 6(5): p. 501–6.

11. Shen, W., et al., Dichotomous dopaminergic control of striatal synaptic plasticity. Science, 2008. 321(5890): p. 848–51.

12. Aldrin-Kirk, P., et al., Chemogenetic modulation of cholinergic interneurons reveals their regulating role on the direct and indirect output pathways from the striatum. Neurobiol Dis, 2018. 109 (Pt A): p. 148–162.

13. Ding, Y., et al., Enhanced striatal cholinergic neuronal activity mediates L-DOPA-induced dyskinesia in parkinsonian mice. Proceedings of the National Academy of Sciences of the United States of America, 2011. 108(2): p. 840–5.

14. Divito, C.B., et al., Loss of VGLUT3 Produces Circadian-Dependent Hyperdopaminergia and Ameliorates Motor Dysfunction and l-Dopa-Mediated Dyskinesias in a Model of Parkinson’s Disease. J Neurosci, 2015. 35(45): p. 14983–99.

15. Gangarossa, G., et al., Role of the atypical vesicular glutamate transporter VGLUT3 in l-DOPA-induced dyskinesia. Neurobiol Dis, 2016. 87: p. 69–79.

16. Lim, S.A.O., et al., Enhanced histamine H2 excitation of striatal cholinergic interneurons in L-DOPA-induced dyskinesia. Neurobiol Dis, 2015. 76: p. 67–76.

17. Won, L., et al., Striatal cholinergic cell ablation attenuates L-DOPA induced dyskinesia in Parkinsonian mice. The Journal of neuroscience: the official journal of the Society for Neuroscience, 2014. 34(8): p. 3090–4.

18. Duvoisin, R.C., Cholinergic-anticholinergic antagonism in parkinsonism. Arch Neurol, 1967. 17(2): p. 124–36.

19. Choi, S.J., et al., Alterations in the intrinsic properties of striatal cholinergic interneurons after dopamine lesion and chronic L-DOPA. Elife, 2020. 9.

20. Tanimura, A., et al., Striatal cholinergic interneurons and Parkinson’s disease. Eur J Neurosci, 2018. 47(10): p. 1148–1158.

21. Lim, S.A., U.J. Kang, and D.S. McGehee, Striatal cholinergic interneuron regulation and circuit effects. Frontiers in synaptic neuroscience, 2014. 6: p. 22.

22. Bordia, T., et al., Optogenetic activation of striatal cholinergic interneurons regulates L-dopa-induced dyskinesias. Neurobiology of disease, 2016. 91: p. 47–58.

23. Shen, W., et al., M4 Muscarinic Receptor Signaling Ameliorates Striatal Plasticity Deficits in Models of L-DOPA-Induced Dyskinesia. Neuron, 2016. 90(5): p. 1139.

24. Dawson, V.L., et al., Evidence for dopamine D-2 receptors on cholinergic interneurons in the rat caudate-putamen. Life Sci, 1988. 42(20): p. 1933–9.

25. Centonze, D., et al., Receptor subtypes involved in the presynaptic and postsynaptic actions of dopamine on striatal interneurons. J Neurosci, 2003. 23(15): p. 6245–54.

26. Yan, Z., W.J. Song, and J. Surmeier, D2 dopamine receptors reduce N-type Ca2+ currents in rat neostriatal cholinergic interneurons through a membrane-delimited, protein-kinase-C-insensitive pathway. J Neurophysiol, 1997. 77(2): p. 1003–15.

27. Cabrera-Vera, T.M., et al., RGS9-2 modulates D2 dopamine receptor-mediated Ca2+ channel inhibition in rat striatal cholinergic interneurons. Proc Natl Acad Sci U S A, 2004. 101(46): p. 16339–44.

28. Carr, D.B., et al., Transmitter modulation of slow, activity-dependent alterations in sodium channel availability endows neurons with a novel form of cellular plasticity. Neuron, 2003. 39(5): p. 793–806.

29. Maurice, N., et al., D2 dopamine receptor-mediated modulation of voltage-dependent Na+ channels reduces autonomous activity in striatal cholinergic interneurons. J Neurosci, 2004. 24(46): p. 10289–301.

30. Deng, P., Y. Zhang, and Z.C. Xu, Involvement of I(h) in dopamine modulation of tonic firing in striatal cholinergic interneurons. J Neurosci, 2007. 27(12): p. 3148–56.

31. Bennett, B.D. and C.J. Wilson, Synaptic regulation of action potential timing in neostriatal cholinergic interneurons. J Neurosci, 1998. 18(20): p. 8539–49.

32. Suzuki, T., et al., Dopamine-dependent synaptic plasticity in the striatal cholinergic interneurons. J Neurosci, 2001. 21(17): p. 6492–501.

33. Oswald, M.J., et al., Potentiation of NMDA receptor-mediated transmission in striatal cholinergic interneurons. Front Cell Neurosci, 2015. 9: p. 116.

34. Castello, J., et al., The Dopamine D5 receptor contributes to activation of cholinergic interneurons during L-DOPA induced dyskinesia. Sci Rep, 2020. 10(1): p. 2542.

35. Chuhma, N., et al., Dopamine neurons control striatal cholinergic neurons via regionally heterogeneous dopamine and glutamate signaling. Neuron, 2014. 81(4): p. 901–12.

36. Hnasko, T.S., et al., Vesicular glutamate transport promotes dopamine storage and glutamate corelease in vivo. Neuron, 2010. 65(5): p. 643–56.

37. Tritsch, N.X., et al., Midbrain dopamine neurons sustain inhibitory transmission using plasma membrane uptake of GABA, not synthesis. Elife, 2014. 3: p. e01936.

38. Gonzalez-Reyes, L.E., et al., Sonic hedgehog maintains cellular and neurochemical homeostasis in the adult nigrostriatal circuit. Neuron, 2012. 75(2): p. 306–19.

39. Traiffort, E., et al., High expression and anterograde axonal transport of aminoterminal sonic hedgehog in the adult hamster brain. The European journal of neuroscience, 2001. 14(5): p. 839–50.

40. Huang, Z. and S. Kunes, Hedgehog, transmitted along retinal axons, triggers neurogenesis in the developing visual centers of the Drosophila brain. Cell, 1996. 86(3): p. 411–22.

41. Tokhunts, R., et al., The full-length unprocessed hedgehog protein is an active signaling molecule. The Journal of biological chemistry, 2010. 285(4): p. 2562–8.

42. Su, Y., et al., High frequency stimulation induces sonic hedgehog release from hippocampal neurons. Sci Rep, 2017. 7: p. 43865.

43. Chuhma, N., et al., Dopamine neuron glutamate cotransmission evokes a delayed excitation in lateral dorsal striatal cholinergic interneurons. Elife, 2018. 7.

44. Mingote, S., et al., Dopamine-glutamate neuron projections to the nucleus accumbens medial shell and behavioral switching. Neurochem Int, 2019. 129: p. 104482.

45. Cai, Y. and C.P. Ford, Dopamine Cells Differentially Regulate Striatal Cholinergic Transmission across Regions through Corelease of Dopamine and Glutamate. Cell Rep, 2018. 25(11): p. 3148–3157 e3.

46. Poulin, J.F., et al., Mapping projections of molecularly defined dopamine neuron subtypes using intersectional genetic approaches. Nat Neurosci, 2018. 21(9): p. 1260–1271.

47. Sebastianutto, I., et al., Validation of an improved scale for rating l-DOPA-induced dyskinesia in the mouse and effects of specific dopamine receptor antagonists. Neurobiology of disease, 2016. 96: p. 156–170.

48. Lundblad, M., et al., A model of L-DOPA-induced dyskinesia in 6-hydroxydopamine lesioned mice: relation to motor and cellular parameters of nigrostriatal function. Neurobiology of disease, 2004. 16(1): p. 110–23.

49. Chen, J.K., et al., Small molecule modulation of Smoothened activity. Proceedings of the National Academy of Sciences of the United States of America, 2002. 99(22): p. 14071–6.

50. Frank-Kamenetsky, M., et al., Small-molecule modulators of Hedgehog signaling: identification and characterization of Smoothened agonists and antagonists. Journal of biology, 2002. 1(2): p. 10.

51. Nunes, I., et al., Pitx3 is required for development of substantia nigra dopaminergic neurons. Proc Natl Acad Sci U S A, 2003. 100(7): p. 4245–50.

52. Hwang, D.Y., et al., Selective loss of dopaminergic neurons in the substantia nigra of Pitx3-deficient aphakia mice. Brain Res Mol Brain Res, 2003. 114(2): p. 123–31.

53. Ding, Y., et al., Chronic 3,4-dihydroxyphenylalanine treatment induces dyskinesia in aphakia mice, a novel genetic model of Parkinson’s disease. Neurobiology of disease, 2007. 27(1): p. 11–23.

54. Sinha, S. and J.K. Chen, Purmorphamine activates the Hedgehog pathway by targeting Smoothened. Nat Chem Biol, 2006. 2(1): p. 29–30.

55. Pahwa, R., et al., Amantadine extended release for levodopa-induced dyskinesia in Parkinson’s disease (EASED Study). Movement disorders: official journal of the Movement Disorder Society, 2015. 30(6): p. 788–95.

56. Stanley, P., et al., Meta-analysis of amantadine efficacy for improving preclinical research reliability. Mov Disord, 2018. 33(10): p. 1555–1557.

57. Fasano, S., et al., Inhibition of Ras-guanine nucleotide-releasing factor 1 (Ras-GRF1) signaling in the striatum reverts motor symptoms associated with L-dopa-induced dyskinesia. Proc Natl Acad Sci U S A, 2010. 107(50): p. 21824–9.

58. Riobo, N.A., et al., Activation of heterotrimeric G proteins by Smoothened. Proc Natl Acad Sci U S A, 2006. 103(33): p. 12607–12.

59. Guo, Q., et al., Whole-brain mapping of inputs to projection neurons and cholinergic interneurons in the dorsal striatum. PloS one, 2015. 10(4): p. e0123381.

60. McKinley, J.W., et al., Dopamine Deficiency Reduces Striatal Cholinergic Interneuron Function in Models of Parkinson’s Disease. Neuron, 2019. 103(6): p. 1056–1072 e6.

61. Vilim, F.S., et al., Peptide cotransmitter release from motorneuron B16 in aplysia californica: costorage, corelease, and functional implications. J Neurosci, 2000. 20(5): p. 2036–42.

62. Whim, M.D. and P.E. Lloyd, Frequency-dependent release of peptide cotransmitters from identified cholinergic motor neurons in Aplysia. Proc Natl Acad Sci U S A, 1989. 86(22): p. 9034–8.

63. Ekstrom, J., et al., Depletion of neuropeptides in rat parotid glands and declining atropine-resistant salivary secretion upon continuous parasympathetic nerve stimulation. Regul Pept, 1985. 11(4): p. 353–9.

64. Matamales, M., et al., Aging-Related Dysfunction of Striatal Cholinergic Interneurons Produces Conflict in Action Selection. Neuron, 2016. 90(2): p. 362–73.

65. Matamales, M., J. Gotz, and J. Bertran-Gonzalez, Quantitative Imaging of Cholinergic Interneurons Reveals a Distinctive Spatial Organization and a Functional Gradient across the Mouse Striatum. PloS one, 2016. 11(6): p. e0157682.

66. Bertran-Gonzalez, J., et al., Striatal cholinergic interneurons display activity-related phosphorylation of ribosomal protein S6. PloS one, 2012. 7(12): p. e53195.

67. Barbeau, A., The pathogenesis of Parkinson’s disease: a new hypothesis. Can Med Assoc J, 1962. 87: p. 802–7.

68. Conti, M.M., N. Chambers, and C. Bishop, A new outlook on cholinergic interneurons in Parkinson’s disease and L-DOPA-induced dyskinesia. Neurosci Biobehav Rev, 2018. 92: p. 67–82.

69. Lehmann, J. and S.Z. Langer, The striatal cholinergic interneuron: synaptic target of dopaminergic terminals? Neuroscience, 1983. 10(4): p. 1105–20.

70. Dessaud, E., A.P. McMahon, and J. Briscoe, Pattern formation in the vertebrate neural tube: a sonic hedgehog morphogen-regulated transcriptional network. Development, 2008. 135(15): p. 2489–503.

71. Yan, Z. and D.J. Surmeier, Muscarinic (m2/m4) receptors reduce N-and P-type Ca2+ currents in rat neostriatal cholinergic interneurons through a fast, membrane-delimited, G-protein pathway. The Journal of neuroscience: the official journal of the Society for Neuroscience, 1996. 16(8): p. 2592–604.

72. Goldberg, J.A. and C.J. Wilson, Control of spontaneous firing patterns by the selective coupling of calcium currents to calcium-activated potassium currents in striatal cholinergic interneurons. J Neurosci, 2005. 25(44): p. 10230–8.

73. Voon, V., et al., Impulse control disorders and levodopa-induced dyskinesias in Parkinson’s disease: an update. Lancet Neurol, 2017. 16(3): p. 238–250.

74. Weintraub, D., et al., Impulse control disorders in Parkinson disease: a cross-sectional study of 3090 patients. Arch Neurol, 2010. 67(5): p. 589–95.

75. Lundblad, M., et al., Pharmacological validation of behavioural measures of akinesia and dyskinesia in a rat model of Parkinson’s disease. Eur J Neurosci, 2002. 15(1): p. 120–32.

76. van den Munckhof, P., et al., Pitx3 is required for motor activity and for survival of a subset of midbrain dopaminergic neurons. Development, 2003. 130(11): p. 2535–42.

## References

1. Ding, Y., et al., Chronic 3,4-Dihydroxyphenylalanine Treatment Induces Dyskinesia in Aphakia Mice, a Novel Genetic Model of Parkinson’s Disease. Neurobiology of disease, 2007. 27(1): p. 11–23.

2. Hwang, D.-Y., et al., Selective loss of dopaminergic neurons in the substantia nigra of Pitx3-deficient aphakia mice. Molecular Brain Research, 2003. 114(2): p. 123–131.

3. Gonzalez-Reyes, L.E., et al., Sonic Hedgehog Maintains Cellular and Neurochemical Homeostasis in the Adult Nigrostriatal Circuit. Neuron, 2012. 75(2): p. 306–319.

4. Machold, R., et al., Sonic hedgehog is required for progenitor cell maintenance in telencephalic stem cell niches. Neuron, 2003. 39(6): p. 937–50.

5. Rossi, J., et al., Melanocortin-4 Receptors Expressed by Cholinergic Neurons Regulate Energy Balance and Glucose Homeostasis. Cell Metabolism, 2011. 13(2): p. 195–204.

6. Jeong, J., et al., Hedgehog signaling in the neural crest cells regulates the patterning and growth of facial primordia. Genes & development, 2004. 18(8): p. 937–51.

7. Harfe, B.D., et al., Evidence for an Expansion-Based Temporal Shh Gradient in Specifying Vertebrate Digit Identities. Cell, 2004. 118(4): p. 517–528.

8. Muzumdar, M.D., et al., A global double-fluorescent Cre reporter mouse. genesis, 2007. 45(9): p. 593–605.

9. Francardo, V., et al., Impact of the lesion procedure on the profiles of motor impairment and molecular responsiveness to L-DOPA in the 6-hydroxydopamine mouse model of Parkinson’s disease. Neurobiology of Disease, 2011. 42(3): p. 327–340.

10. Lundblad, M., et al., Pharmacological validation of behavioural measures of akinesia and dyskinesia in a rat model of Parkinson’s disease. European Journal of Neuroscience, 2002. 15(1): p. 120–132.

11. Cenci, M.A. and M. Lundblad, Ratings of L-DOPA-Induced Dyskinesia in the Unilateral 6-OHDA Lesion Model of Parkinson’s Disease in Rats and Mice. Current Protocols in Neuroscience, 2007. 41(1): p. 9.25.1-9.25.23.

12. Sebastianutto, I., et al., Validation of an improved scale for rating l-DOPA-induced dyskinesia in the mouse and effects of specific dopamine receptor antagonists. Neurobiology of disease, 2016. 96: p.156–170.

13. Aubert, I., et al., Increased D1 dopamine receptor signaling in levodopa-induced dyskinesia. Ann Neurol, 2005. 57(1): p. 17–26.

14. Berton, O., et al., Striatal overexpression of DeltaJunD resets L-DOPA-induced dyskinesia in a primate model of Parkinson disease. Biol Psychiatry, 2009. 66(6): p. 554–61.

15. Fox, S.H., et al., A critique of available scales and presentation of the Non-Human Primate Dyskinesia Rating Scale. Mov Disord, 2012. 27(11): p. 1373–8.

16. Sparta, D.R., et al., Construction of implantable optical fibers for long-term optogenetic manipulation of neural circuits. Nature Protocols, 2012. 7(1): p. 12–23.

17. Alcacer, C., et al., Gαolf Mutation Allows Parsing the Role of cAMP-Dependent and Extracellular Signal-Regulated Kinase-Dependent Signaling in L-3,4-Dihydroxyphenylalanine-Induced Dyskinesia. The Journal of Neuroscience, 2012. 32(17): p. 5900–5910.

18. Bertran-Gonzalez, J., et al., Striatal Cholinergic Interneurons Display Activity-Related Phosphorylation of Ribosomal Protein S6. PLOS ONE, 2012. 7(12): p. e53195.

